# Gene-Specific Nonsense-Mediated mRNA Decay Targeting for Cystic Fibrosis Therapy

**DOI:** 10.1101/2021.07.13.452144

**Authors:** Young Jin Kim, Tomoki Nomakuchi, Foteini Papaleonidopoulou, Adrian R. Krainer

## Abstract

Low *CFTR* mRNA expression due to nonsense-mediated mRNA decay (NMD) is a major hurdle in developing a therapy for cystic fibrosis (CF) caused by the *W1282X* mutation in the *CFTR* gene. CFTR-W1282X truncated protein retains partial function, so increasing its levels by inhibiting NMD of its mRNA will likely be beneficial. Because NMD regulates the normal expression of many genes, gene-specific stabilization of *CFTR-W1282X* mRNA expression is more desirable than general NMD inhibition. Synthetic antisense oligonucleotides (ASOs) designed to prevent binding of exon junction complexes (EJC) downstream of premature termination codons (PTCs) attenuate NMD in a gene-specific manner. We developed a cocktail of three ASOs that specifically increases the expression of *CFTR* W1282X mRNA and CFTR protein in ASO-transfected human bronchial epithelial cells. This treatment increased the CFTR-mediated chloride current. These results set the stage for clinical development of an allele-specific therapy for CF caused by the W1282X mutation.

## Introduction

*CFTR-W1282X*, the 6^th^ most common CF-causing mutation, causes a severe form of CF and is present in 1.2% of CF patients worldwide ^1^, 2.2% of U.S. CF patients ^2^, and up to 40% of Israeli CF patients ^3^. The CFTR-W1282X truncated protein retains partial function ^4–6^, but is expressed at a very low level, due to nonsense-mediated mRNA decay (NMD). In general, NMD prevents the accumulation of potentially harmful truncated proteins translated from premature termination codon (PTC)-containing mRNAs. However, when NMD reduces the expression of a mutant CFTR protein that has partial activity, it exacerbates the phenotype, so that patients homozygous for the *CFTR-W1282X* mutation or compound heterozygous for *CFTR-W1282X* and another CF-causing mutation have poor clinical outcomes. As low as 10% of normal CFTR function provides a significant therapeutic benefit for CF patients ^7,8^. Thus, increasing the expression of mutant CFTR protein with residual activity is expected to be beneficial.

NMD is a major hurdle for developing a targeted therapy for CF caused by the *CFTR-W1282X* mutation. The approval of CFTR correctors that enhance post-translational CFTR processing, and potentiators that improve CFTR channel opening, brought benefit to the majority of CF patients ^9^. However, these therapeutic options are not effective against CF caused by *CFTR-W1282X*, due to the low expression of *CFTR-W1282X* mRNA. One approach to treat CF caused by this mutation involves read-through compounds (RTCs) that increase the level of full-length protein by reducing the fidelity of the ribosome at the PTC ^10^. Gentamicin is a type of RTC that can increase full-length CFTR protein *in vitro*, but its clinical efficacy for various CF nonsense mutations is limited by NMD ^11,12^. Likewise, ataluren is another non-aminoglycoside RTC with a very good safety profile, but it did not improve forced expiratory volume (FEV) in CF patients with various nonsense mutations, including *W1282X*, in clinical trials ^13,14^.

Preclinical models of CF caused by the *W1282X* mutation showed that the efficacy of RTCs can be increased *in vivo* by knocking down key components of the NMD pathway ^15^. However, a clinically viable NMD-suppression approach does not exist yet. CFTR potentiators and correctors such as ivacaftor (VX-770) and lumacaftor (VX-809) enhance CFTR-W1282X activity *in vitro*, and may potentially benefit patients with the W1282X mutation, but they are not effective if the truncated protein expression is too low ^4–6,16^. Thus, there is a pressing need for strategies to overcome NMD of the *CFTR-W1282X* mRNA.

Several NMD-suppression strategies have been developed for potential application to diseases caused by NMD-sensitive nonsense mutations. These include inhibition of NMD by small-molecule inhibitors ^17–19^ or knockdown of key NMD factors ^20, 19^. However, global inhibition of NMD may be detrimental, because the NMD machinery targets a subset of normal and physiologically functional mRNA isoforms, thereby post-transcriptionally regulating gene expression ^21^. Therefore, global inhibition of NMD could disrupt mRNA homeostasis in a broad range of tissues ^15^.

NMD is strongly dependent on a complex of RNA-binding proteins called the exon junction complex (EJC). In contrast to many RNA-binding proteins, EJCs bind mRNA in a position-dependent, sequence-independent manner ^22–25^. More than 80% of EJCs are positioned 20∼24 nucleotides (nt) upstream of an exon-exon junction ^22–26^. Normal stop codons are typically in the last exon, whereas many PTCs are upstream of one or more exon-exon junctions, and thus are upstream of at least one EJC ^27^. The ‘55-nt rule’ predicts that mRNAs with a PTC > 55 nt upstream of the last exon-exon junction are degraded by NMD, reflecting the footprint of the stalled ribosome ^23^. A downstream EJC interacts with the ribosome stalled at the PTC, and recruits NMD factors to form a degradation complex that promotes decapping, deadenylation, and endocleavage of the target mRNA, which is subsequently degraded ^28^. Thus, the EJC is a major enhancer of NMD.

Disrupting the downstream EJC association with PTC-containing mRNA can be used to inhibit NMD ^29^. Uniformly 2’-*O*-(2-methoxyethyl) (MOE)-modified antisense oligonucleotides (ASOs) can stably hybridize to complementary RNAs without triggering RNase H-mediated degradation ^30^. Such ASOs are effective tools for disrupting the interaction between an RNA and its binding proteins, and can be used to alter mRNA processing and translation *in vitro* and *in vivo* ^30–34^. We previously developed ASOs that target presumptive downstream EJC sites of PTC-containing mRNAs. These ASOs efficiently attenuate NMD of their target genes, restoring mRNA and proteins levels ^29^. In the present study, we demonstrate that a cocktail of three ASOs targeting presumptive downstream EJC binding sites specifically increases the expression of endogenous *CFTR*-*W1282X* mRNA in human bronchial epithelial (HBE) cells. Furthermore, the ASO cocktail increases partially active CFTR protein and CFTR-mediated chloride current in HBE cells. These results set the stage for the clinical development of an allele-specific therapy for CF caused by the *W1282X* mutation.

## Results

*CFTR-W1282X* mRNA has four downstream exon-exon junctions, on exons 23-24, 24-25, 25-26, and 26-27. However, the predicted EJC binding site on exon 23 is approximately 5 nt downstream of the PTC—within the ribosome footprint—and thus only three exons (24, 25, and 26) are predicted to harbor EJCs that can induce NMD. To investigate the impact of each EJC on NMD of *CFTR-W1282X* mRNA, we generated U2OS cells stably expressing doxycycline-inducible *CFTR*-minigene NMD reporters, each with only one presumptive downstream EJC site (Figure 1A, Supplementary Fig.1A). These NMD reporters are three-exon minigenes downstream of GFP, and comprise *CFTR* cDNA sequence for exons 22-27 and intervening sequences (IVSs) that are shortened natural *CFTR* introns.

**Figure 1.**
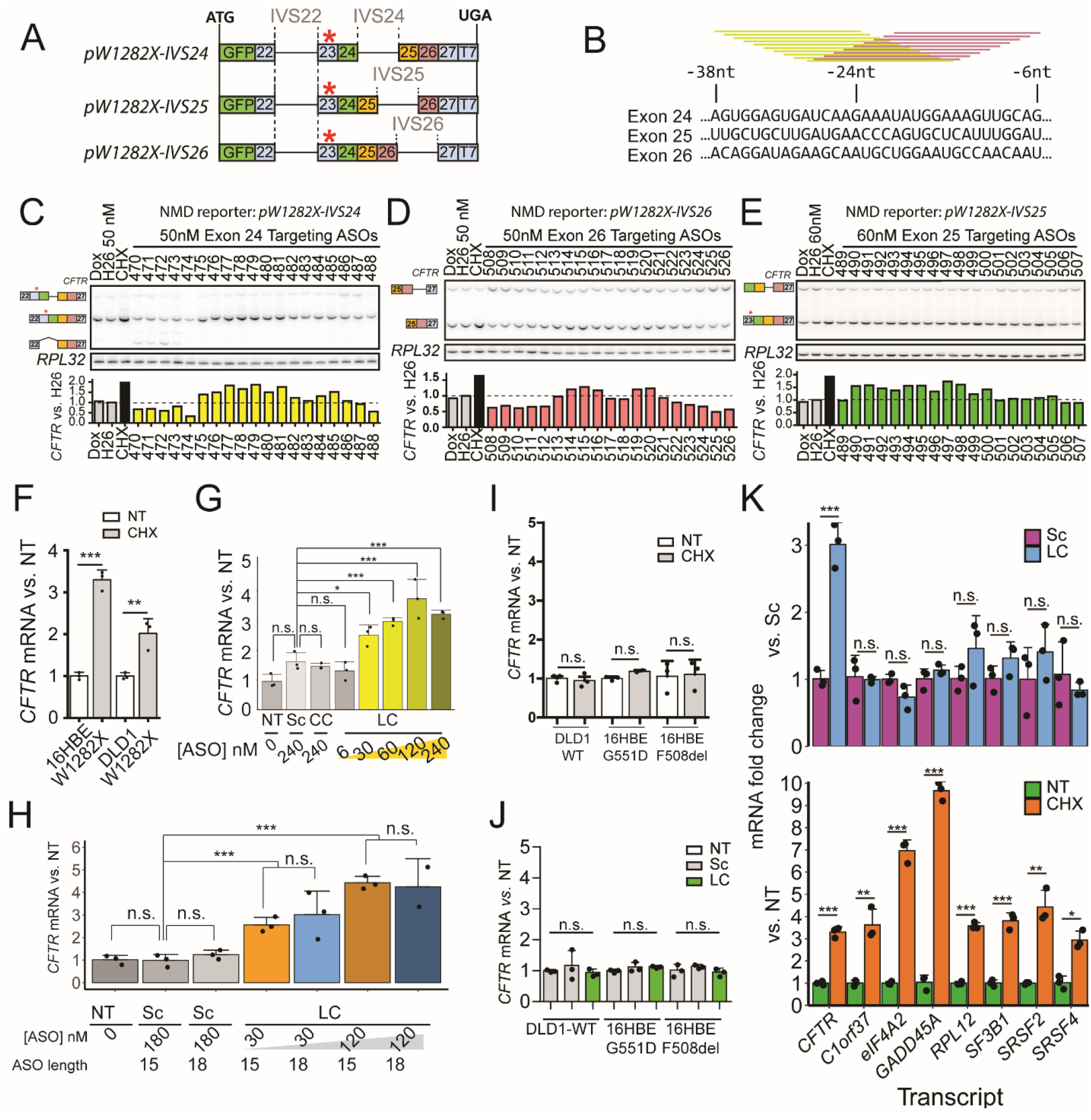
Identification of NMD-inhibiting ASOs and assessment of their specificity. **A**. Schematic of NMD reporters. The numbers show the *CFTR* exons present in the NMD reporters. The red asterisk (*) indicates the location of the W1282X mutation. **B**. Schematic of ASO screening. 19 MOE-PS modified 15mer ASOs (yellow and magenta bars) were designed to cover the presumptive EJC binding sites on exons 24, 25, and 26 at 1-nt resolution. **C-E**. U2OS cells stably expressing each NMD reporter were transfected with individual ASOs targeting EJC binding regions on *CFTR* (C) exon 24, (D) exon 25, or (E) exon 26, respectively. Reporter mRNA levels were measured by radioactive RT-PCR, using primers listed in Table 4. *RPL32-*normalized reporter expression is compared to that of the negative-control ASO (H26) transfection, and is shown below the RT-PCR images. **F**. Effect of cycloheximide (CHX) on *CFTR* expression in 16HBE-W1282X and DLD1-W1282X cells. **G**. Effect of the ASO cocktail C478-C495-C515 (LC) on *CFTR* expression in 16HBE-W1282X cells. **H**. Comparison between 15mer and 18mer lead ASO cocktails (C478-C494-C514 or C24-C25-C26, respectively). The 15mer and 18mer scramble ASOs were used as negative controls. **I**. Effect of cycloheximide (CHX) on *CFTR* expression in DLD1-WT, 16HBE-G551D, and 16HBE-F508del cells. **J**. *CFTR* mRNA levels in DLD1-WT, 16HBE-G551D, and 16HBE-F508del transfected with the control ASOs or the lead ASO cocktail at a nominal total concentration of 120 nM. **K**. Endogenous NMD-sensitive mRNA levels in 16HBE-W1282X cells treated with cycloheximide (orange), 120 nM scramble ASO (Sc; purple) or 120 nM lead ASO cocktail (blue). All mRNA levels in F-K were measured by RT-qPCR. *RPL32* served as internal reference for all panels except panel H, in which *HPRT* served as internal reference. NT=No treatment; Dox: doxycycline 1 μg/mL; Sc=Scramble ASO; CC=Control ASO cocktail C488-C507-C526; LC= lead ASO cocktail C478-C495-C515, C478-C494-C514, or C24-C25-C26; CHX = 1-hr incubation with 100 μg/mL cycloheximide. All error bars indicate standard deviation. For all treatments, n=3, except n=2 in LC18-mer 120nM in panel G. For all statistical tests, n.s. P>0.05, *P<0.05, **P<0.01, ***P<0.001. Panel F, I, and K: Student’s t-test. Panel G, H, and J: one-way ANOVA with Tukey’s post-test.

The reporters showed some intron retention, and *pW1282X-IVS23* generated an additional isoform by use of an alternative 3’ splice site (3’ss; based on size and motif predictions) (Supplementary Fig.1B-F). The ‘55-nt rule’ predicts that an exon-exon junction <55 nt downstream of the PTC does not induce NMD ^23^. As mentioned above, only the EJCs on exon 24, 25, and 26 are predicted to induce NMD. Inhibiting NMD with cycloheximide increased the mRNA levels of the reporters harboring the PTC and intron 24, 25, or 26 (*pW1282X-IVS24, pW1282X-IVS25*, or *pW1282X-IVS26*) (Supplementary Fig.1D-F); conversely, the reporters harboring the PTC and intron 23 (*pW1282X-IVS23*) or harboring intron 24 but not the PTC (*pWT-IVS24*) were not sensitive to cycloheximide (Supplementary Fig.1B-C).

Uniformly 2’-*O*-(2-methoxyethyl) (MOE)-modified ASOs can stably hybridize to complementary mRNAs without inducing RNAse-H-mediated degradation, and modulate their posttranscriptional processing, including NMD ^31^. Using our previously described ASO screening strategy for gene-specific antisense inhibition of NMD--dubbed “GAIN” ^29^—we designed sets of 19 overlapping 15mer ASOs to target each of the presumptive EJC binding sites on the NMD reporters *pW1282X-IVS24, pW1282X-IVS25*, and *pW1282X-IVS26*, respectively (Fig. 1B). We screened a total of 57 ASOs, uniformly modified with MOE ribose and a phosphorothioate (P=S) backbone. ASOs H24, H26, and M33 are uniformly MOE and P=S modified negative-control ASOs that are not complementary to any gene expressed in the reporter-expressing cells ^29^. Based on the screen, we chose C478 and C515 as the initial lead ASOs targeting exons 24 and 26, respectively (Fig. 1C-D). Because the retention of IVS25 in *pW1282X-IVS25* caused by some ASOs prevented a clear assessment of NMD inhibition by the screened ASOs (Supplementary Fig. 2A), we generated a new *pW1282X-IVS25* NMD reporter with a stronger 5’ splice site (5’ss) but the same amino acid sequence (Supplementary Fig. 2B). Among several ASOs that increased the new NMD reporter levels, we chose C495 as the lead ASO (Fig. 1E). The candidate ASOs inhibited NMD of the reporters in a dose-dependent manner (Supplementary Fig. 2C-H).

Endogenous *CFTR* mRNA is targeted for NMD in human bronchial epithelial cells and colon cancer cells harboring the homozygous *CFTR-W1282X* mutation (16HBE-W1282X and DLD1-W1282X cells) (Fig. 1F). We used two negative-control treatments: i) a scrambled-sequence ASO based on C494; and ii) a cocktail composed of C488+C507+C526 ASOs, which did not stabilize the NMD reporters. We first tested a lead GAIN ASO cocktail composed of C478+C495+C515, based on the above NMD-reporter screening results. Compared to the negative-control ASOs, the C478+C495+C515 ASO cocktail significantly increased *CFTR* mRNA levels in both 16HBE-W1282X and DLD1-W1282X cells (Fig. 1G, Supplementary Fig. 3).

Length is an important parameter in ASO design that can affect the efficacy and specificity of uniformly modified ASOs ^35–37^. Based on the results of the above 15mer ASO screens, we designed a new 18mer-ASO cocktail and an 18mer scramble-ASO control. Transfection of the scramble ASO did not increase *CFTR-W1282X* mRNA levels in 16HBE-W1282X cells, whereas the 18mer ASO cocktail C24+C25+C26 increased *CFTR-W1282X* mRNA levels in a dose-dependent manner (Fig. 1H). The 18mer ASO cocktail did not affect two other endogenous NMD-sensitive mRNAs, *eIF4A2* and *SRSF2* (Supplementary Fig. 4). Thus, the lead GAIN ASO 18mer cocktail inhibited NMD of *CFTR*-*W1282X* mRNA in a gene-specific manner.

Based on the presumptive mechanism of action of the ASO cocktails, they should not affect the total mRNA levels of wild-type (WT) or missense-mutant *CFTR* mRNAs that are not sensitive to NMD (Fig. 1I). Indeed, *CFTR* mRNA levels were insensitive to transfection of the control ASO or C478+C495+C515 ASO cocktail in DLD1-WT, 16HBE-F508del, and 16HBE-G551D cells (Fig. 1J). Thus, the C478+C495+C515 ASO cocktail allele-specifically increased the mRNA levels of nonsense-mutant *CFTR-W1282X* mRNA. To test whether C478+C495+C515 inhibits NMD of *CFTR-W1282X* mRNA specifically, as opposed to somehow affecting global NMD, we used RT-qPCR to survey seven other endogenous NMD-sensitive transcripts that are upregulated upon NMD inhibition by cycloheximide treatment ^38,39^. As expected, C478+C495+C515 increased *CFTR-W1282X* mRNA, without significantly changing any of the other NMD-sensitive mRNAs (Fig. 1K).

To identify the optimal GAIN ASO cocktail, we performed a more comprehensive screening in 16HBE-W1282X cells (Fig. 2A-C). To systematically screen ASOs targeting each exon, we tested cocktails composed of two constant ASOs and one varying ASO. For example, to screen exon-24-targeting ASOs, we tested 19 cocktails composed of varying exon-24-targeting ASOs and the same two ASOs targeting exons 25 and 26. After testing 57 such ASO cocktails, we identified C478+C494+C514 as the new lead GAIN ASO cocktail, which was more potent than C478+C495+C515 (Fig. 2D). At the highest concentration tested, both ASO cocktails increased *CFTR-W1282X* mRNA similarly, but at a lower concentration, the new lead cocktail had significantly higher potency.

**Figure 2.**
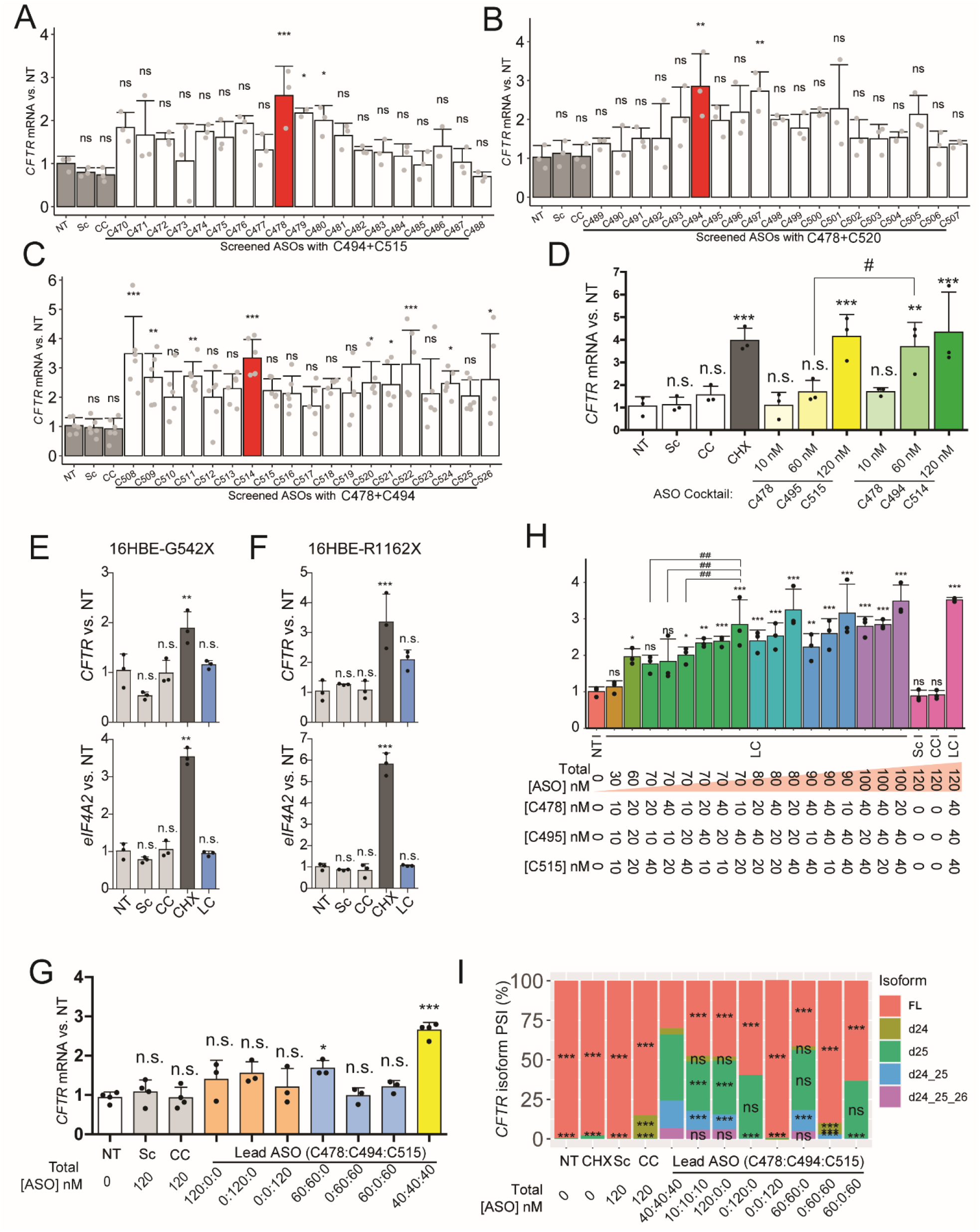
ASO-cocktail optimization and mechanism of action. **A-C**. New ASOs targeting *CFTR* (H) exon 24, (I) exon 25, or (J) exon 26 were individually screened in 16HBE-W1282X cells, in combination with two ASOs that target the other two exons, at a total nominal concentration of 120 nM. Red bars indicate the lead ASO identified in each combination screen. **D**. Comparison between the ASO cocktails C478-C495-C515 and C478-C494-C514. **E-F**. *CFTR* and *eIF4A2* mRNA levels in (D) 16HBE-G542X cells and (E) 16HBE-R1162X cells transfected with ASOs at a total nominal concentration of 120 nM. **G**. The number of required EJCs targeted by ASOs C478, C494, C515, or all together, was assessed by transfecting 16HBE-W1282X cells with one, two, or three EJC-targeting ASOs at the same total nominal concentration. **H**. 16HBE-W1282X cells were transfected with various combinations of the lead ASOs C478, C495, and C515. **I**. Mean PSI of each *CFTR* isoform in 16HBE-W1282X cells transfected with various combinations of C478, C495, and C515. All mRNA levels in A-G were measured by RT-qPCR. *RPL32* mRNA level served as an internal reference. Error bars show standard deviation. Abbreviations are as in Figure 1. n=3 for all treatments, except in panel C, n=5 or 6, and panel G, n=4 for NT, Sc, CC, and Lead ASO 40:40:40. For all Dunnett’s post-tests, n.s. P>0.05, *P<0.05, **P<0.01, ***P<0.001. For all Student’s t-tests, #P<0.05, ##P<0.01. Panels A-H: one-way ANOVA with Dunnett’s post-test versus NT, or Student’s t-test. Panel I: n=4, one-way ANOVA with Dunnett’s post-test, versus each isoform in ‘lead ASO 40:40:40’.

*G542X* and *R1162X CFTR* mutations are on exons 12 and 22, respectively. As these mutant mRNAs harbor more than three EJCs downstream of the premature termination codon, we tested whether their levels would be insensitive to the lead ASO cocktail C478+C494+C514. We transfected 16HBE-G542X and 16HBE-R1162X cells harboring homozygous *G542X* and *R1162X* mutations, respectively, with control ASOs or the lead ASO cocktail C478+C494+C514 (Fig. 2E and F). Compared to the no-treatment control, transfection of 120 nM scramble ASO, control ASO cocktail, or C478+C494+C514 did not affect the levels of *CFTR-G542X, CFTR-R1162X*, and NMD-sensitive *eIF4A2* mRNAs. Only cycloheximide treatment caused significant increases in these NMD-sensitive mRNA levels. These results are consistent with the EJC-centric model of NMD, according to which at least one EJC >55nt downstream of a PTC is sufficient to induce strong NMD ^23^.

Using various combinations of the NMD-inhibiting ASOs, we next tested whether all three presumptive downstream EJC binding sites on *CFTR-W1282X* mRNA must be targeted with the corresponding ASOs for effective mRNA stabilization. Targeting only one or two EJC binding sites with the respective lead ASOs partially stabilized the *CFTR-W1282X* mRNA, but the most significant and efficient increase in *CFTR-W1282X* mRNA was obtained by simultaneous transfection of all three lead ASOs, in both 16HBE-W1282X and DLD1-W1282X cells (Fig. 2G, Supplementary Fig. 5). Because ASO cocktails composed of one or two ASOs elicited a small increase in *CFTR-W1282X* mRNA levels, we next asked whether certain presumptive EJC binding sites may be more important than others. To test this possibility, we transfected 16HBE-W1282X cells with ASO cocktails with varying ratios of the individual ASOs (Fig. 2H). The C478+C495+C515 cocktail with the highest total ASO concentration and an equimolar ratio of 40:40:40 nM increased *CFTR-W1282X* mRNA the most, and the increase was dependent on the total ASO concentration. In general, limiting the concentration of C495 in the cocktail reduced the *CFTR-W1282X* mRNA levels to the greatest extent (Supplementary Fig. 6A-C). Interestingly, ASO cocktails with equal total concentrations, but different ASO ratios, did not have equivalent effects on *CFTR-W1282X* mRNA levels. Also, some ASO cocktails increased *CFTR-W1282X* mRNA similarly or more than others, despite their lower total ASO concentration. The differences among *CFTR-W1282X* mRNA expression changes caused by the various ASO cocktails may be attributable to various factors, including partial EJC occupancy on different exons, differences in ASO uptake and target accessibility or affinity, involvement of EJC-independent NMD pathways, and RNA secondary structure ^40,41^.

Some EJC-targeting ASOs may affect splicing, if their binding site overlaps with cis-elements that regulate splicing. We monitored exon 24-26 splicing by RT-PCR in DLD1-WT cells transfected with ASOs (Supplementary Fig. 7A). Exon-26-targeting ASOs did not detectably disrupt *CFTR* mRNA splicing. On the other hand, all exon-25-targeting ASOs caused slight exon-25 skipping, and some exon-24-targeting ASOs caused substantial exon-24 skipping. As disrupted binding of serine-rich (SR) proteins to exonic splicing enhancers (ESEs) by uniformly modified MOEPS ASOs can cause exon skipping ^42–45^, we used ESEfinder ^46^ to identify putative ESEs on *CFTR* exons 24-26 that might be blocked by the lead ASOs (Supplementary Fig. 7B). mRNA sequences complementary to the lead ASOs C478 and C494 overlap with SR protein motifs; however, overlap with an SR protein motif is insufficient to predict an ASO’s interference with splicing.

Transfection of the 15-mer or 18-mer lead ASOs caused dose-dependent, multiple exon skipping in human bronchial cells (Fig. 2I, Supplementary Fig. 8A-F): single (exon 24 and 25 skipping: d24 and d25), double (exon 24-25 skipping: d24-25), and triple exon skipping (exon 24-25-26 skipping: d24-25-26) of *CFTR* mRNA. Similar splicing changes occurred with the 15mer or 18mer lead ASO cocktail treatment by free uptake (Supplementary Fig. 9A-B). The splicing changes were not cell-line-specific, as the lead ASO cocktail promoted similar splicing changes in 16HBE-W1282X and DLD-W1282X cells (Supplementary Fig. 10A-B).

Control ASO cocktails caused a smaller degree of exon 24 and 25 skipping, consistent with the results in DLD1-WT cells (Fig. 2I). The 15mer and 18mer lead ASO cocktails generated the same *CFTR* isoforms, but with varying degrees of percent-spliced-in (PSI); for example, the PSI of the exon 24-25 double-skipping event was higher in 16HBE-W1282X cells treated with the 18mer lead cocktail (Supplementary Fig. 8C-D and 9B). Interestingly, whereas the 15mer or 18mer exon-26-targeting ASOs alone did not cause exon 26 skipping, the ASO cocktails containing exon-24-targeting ASO caused the appearance of the triple-skipped isoform (d24-25-26) (Fig. 2I, Supplementary Fig. 8C-D). To search for potential off-target sites on *CFTR* pre-mRNA, we looked for sites complementary to C478 with a maximum of four nucleotide mismatches, downstream of exon 24, and found only one site with four mismatches in intron 23. The chance of finding an off-target with ≥ 4-nt mismatches is very low ^47^, suggesting that C478 is unlikely to cause exon 25 and 26 skipping by binding to ESEs in these exons. These results suggest that an ESE and/or the EJC in exon 24 is involved in long-range splicing regulation. Recent studies showed that EJCs help maintain faithful splicing transcriptome-wide ^48–54^.

Despite the splicing alterations, the increase in the total *CFTR* mRNA by the lead ASO cocktail resulted in increased CFTR-W1282X protein levels, compared to the scramble ASO control (Fig. 3A-B). This result was expected, because none of the splicing changes affect the reading frame upstream of the nonsense mutation in exon 23. Combining the lead ASO cocktail with lumacaftor (VX-809), a corrector that improves the folding of CFTR protein ^5,55^, further increased the total CFTR-W1282X protein levels (Fig. 3A-B). Three different patterns of CFTR bands are visible on a Western blot: non-glycosylated A-band, core glycosylated B-band, and fully mature, glycosylated C-band ^56^. As shown previously ^5^, truncated CFTR-W1282X exists as core-glycosylated and mature glycosylated forms (Supplementary Fig. 11A). Extensive enzymatic deglycosylation revealed that truncated-core and fully-mature-glycosylated CFTR-W1282X proteins are upregulated by transfection of the lead ASO cocktail in 16HBE-W1282X cells (Supplementary Fig. 11B and C).

**Figure 3.**
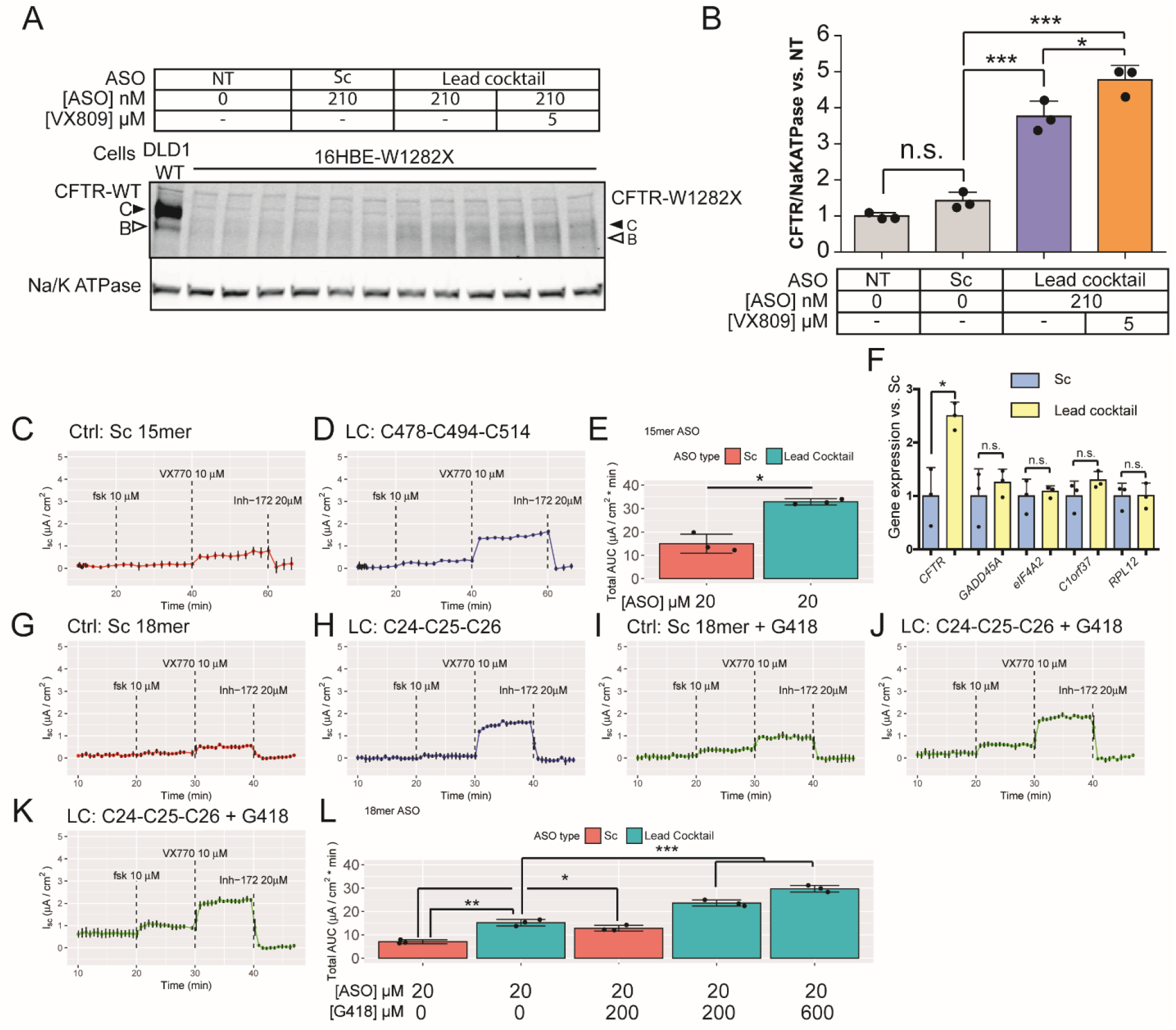
Effect of the lead ASO cocktail on CFTR-W1282X protein expression and function. **A**. Western blot of CFTR-W1282X protein in 16HBE-W1282X cells transfected with control or EJC-targeting ASO cocktail C478-494-C515 and treated with VX-809 at the indicated concentrations. The closed and open arrowheads indicate C and B bands of CFTR proteins, respectively. **B**. Quantification of total CFTR protein in (A). **C-D**. Average traces from Ussing-chamber assay of 16HBE-W1282X cells treated with 20 μM of (C) 15mer scramble ASO (Sc) or (D) 15mer lead ASO cocktail C479-C494-C514. **E**. Total area under the curve in (C) and (D). Before the assays, the cells were treated for 24 h with VX-809 (3 μM). The traces shown are the average of the three replicates. **F**. The levels of *CFTR-W1282X* and endogenous NMD-sensitive mRNAs in 16HBE-W1282X cells assayed in €, normalized to Sc control. *RPL32* was used as internal reference. **G-K** Average traces from Ussing-chamber assay of 16HBE-W1282X cells treated with (G) 20 μM 18mer scramble ASO, (H) 20 μM 18mer lead ASO cocktail (C24-C25-C26), (I) 20 μM 18mer scramble ASO and 200 μM G418, (J) 20 μM 18mer lead ASO cocktail and 200 μM G418, and (K) 20 μM 18mer lead ASO cocktail and 600 μM G418. All G418 treatments were started 24 hr prior to the assay. The traces shown are the average of three replicates. Black error bars on each point show standard deviation. **L**. The total area under the curve of (G-K). Error bars show standard deviations. Fsk: forskolin. n=3 for all treatments. For all statistical tests, n.s. P>0.05, *P<0.05, **P<0.01, ***P<0.001. Panels E and F: Student’s t-test. Panels B and L: one-way ANOVA with Tukey’s post-test.

We next measured CFTR function in 16HBE-W1282X cells treated with the lead GAIN ASO cocktail, with the Ussing-chamber assay ^57^. FDA-approved CFTR potentiators and correctors, such as ivacaftor (VX-770) and lumacaftor (VX-809), can enhance CFTR-W1282X activity *in vitro*, and may potentially benefit patients with the W1282X mutation ^4–6,16^. Thus, we combined all ASO treatments for Ussing-chamber assays with VX-809 and VX-770. The 16HBE-W1282X cells treated with the lead 15mer ASO cocktail (C478+C494+C514) showed increased CFTR-mediated chloride current, compared to 16HBE-W1282X cells treated with scramble 15mer ASO (Fig. 3C-E). We verified that the 16HBE-W1282X cells transfected with the lead ASO cocktail while cultured on transwell plates for the Ussing chamber assay showed gene-specific increase in *CFTR-W1282X* mRNA levels (Fig. 3F). Similar to the 15mer lead ASO cocktail, the 18mer lead ASO cocktail (C24+C25+C26) significantly increased CFTR function, compared to the control 18mer scramble ASO treatment (Fig. 3G-L). This result demonstrates for the first time that gene-specific NMD inhibition of a hypomorphic *CFTR* allele leads to an increase in CFTR-mediated chloride current. All *CFTR-W1282X* mRNA isoforms generated by the ASO cocktail treatment presumably terminate at the *W1282X* codon, but the contribution of each isoform to CFTR activity may be affected by various factors, including mRNA stability, transport to the cytoplasm, and translational efficiency.G418 is an aminoglycoside antibiotic that at high concentrations induces translational read-through of reporters containing *CFTR* nonsense mutations, and increases truncated CFTR-W1282X protein levels by NMD inhibition ^18,58,59^. Indeed, 400 μM (0.2 mg/mL) G418 alone increased truncated CFTR-W1282X protein levels (Supplementary Fig. 11D-E), and 200 μM (0.1 mg/mL) increased CFTR activity in 16HBE-W1282X cells (Fig. 3I and L). However, full-length CFTR reflecting read-through activity remained below the level of detection by Western blotting (Supplementary Fig. 11C-D). Combining 200 μM or 600 μM G418 with the lead ASO cocktail further increased CFTR-W1282X function, compared to the respective lead ASO cocktail treatment alone (Fig. 3J-L). This result suggests that some level of read-through at the *W1282X* codon may occur.

## Discussion

NMD severely limits the therapeutic development for CF caused by *CFTR-W1282X* mutation. Global NMD suppression can be achieved by targeting key NMD factors by gap-mer ASOs that induce gene knockdown or small molecules that inhibit the activity of NMD factors ^17–20^. However, targeted NMD suppression may be more desirable as it can avoid unwanted side-effects that can be caused by non-specific NMD inhibition. Here, we demonstrate for the first time that gene- and allele-specific NMD suppression using EJC-targeting ASO cocktails increases truncated CFTR-W1282X protein, as well as CFTR function.

The lead GAIN ASO cocktails inhibited NMD of *CFTR-W1282X* mRNA by targeting presumptive EJC binding sites downstream of the PTC, independently of cell type. Consistent with the EJC-centric model of NMD ^40^, we achieved efficient NMD suppression only when all downstream EJC binding sites that contribute to NMD (i.e., those on exons 24-26) were targeted by ASOs. The lead ASO cocktails did not inhibit NMD of other endogenous NMD-sensitive transcripts we tested, or affect the levels of NMD-insensitive *CFTR* mRNA. Likewise, the lead ASO cocktails did not affect *CFTR* mRNA levels in cells harboring PTCs upstream of exon 23.

Thus, our results demonstrate that the lead ASO cocktails inhibit NMD of *CFTR-W1282X* mRNA by preventing the binding of EJCs located downstream of the PTC and beyond the footprint of the stalled ribosome. These observations rule out the possibility of global NMD suppression due to ASO treatment, or *CFTR* mRNA stabilization by inhibition of other mRNA-degradation pathways.

The lead 15-mer and 18-mer cocktails achieved similar levels of gene-specific NMD suppression, but had different effects on splicing. Compared to 12-mer ASOs, 18-mer ASOs tend to have fewer off-target effects on splicing ^60^, but whether our lead 18-mer ASO cocktail meaningfully reduces off-target effects and toxicity will require further investigation. We conclude that rationally designed ASOs can modulate clinically relevant NMD, expanding the current RNA and oligonucleotide therapeutics toolbox.

## Methods

### ASOs

All ASOs were uniformly modified with 2’-*O*-(2-methoxyethyl) (MOE) ribose, phosphorothioate (P=S) linkages, and 5’-methylcytosine. The 15mer ASOs were obtained from Ionis Pharmaceuticals (Carlsbad, CA) and Integrated DNA Technologies (Coralville, IA), and 18mer ASOs were obtained from Bio-Synthesis (Lewisville, TX). All ASOs were dissolved in water and stored at -20 °C. Stock ASO concentrations were calculated based on the A260 measurement and each ASO’s extinction coefficient *e* (mM^-1^ x cm^-1^ @ 260 nm). The sequences of all ASOs used in this study are listed in Table 1.

**Table 1.**
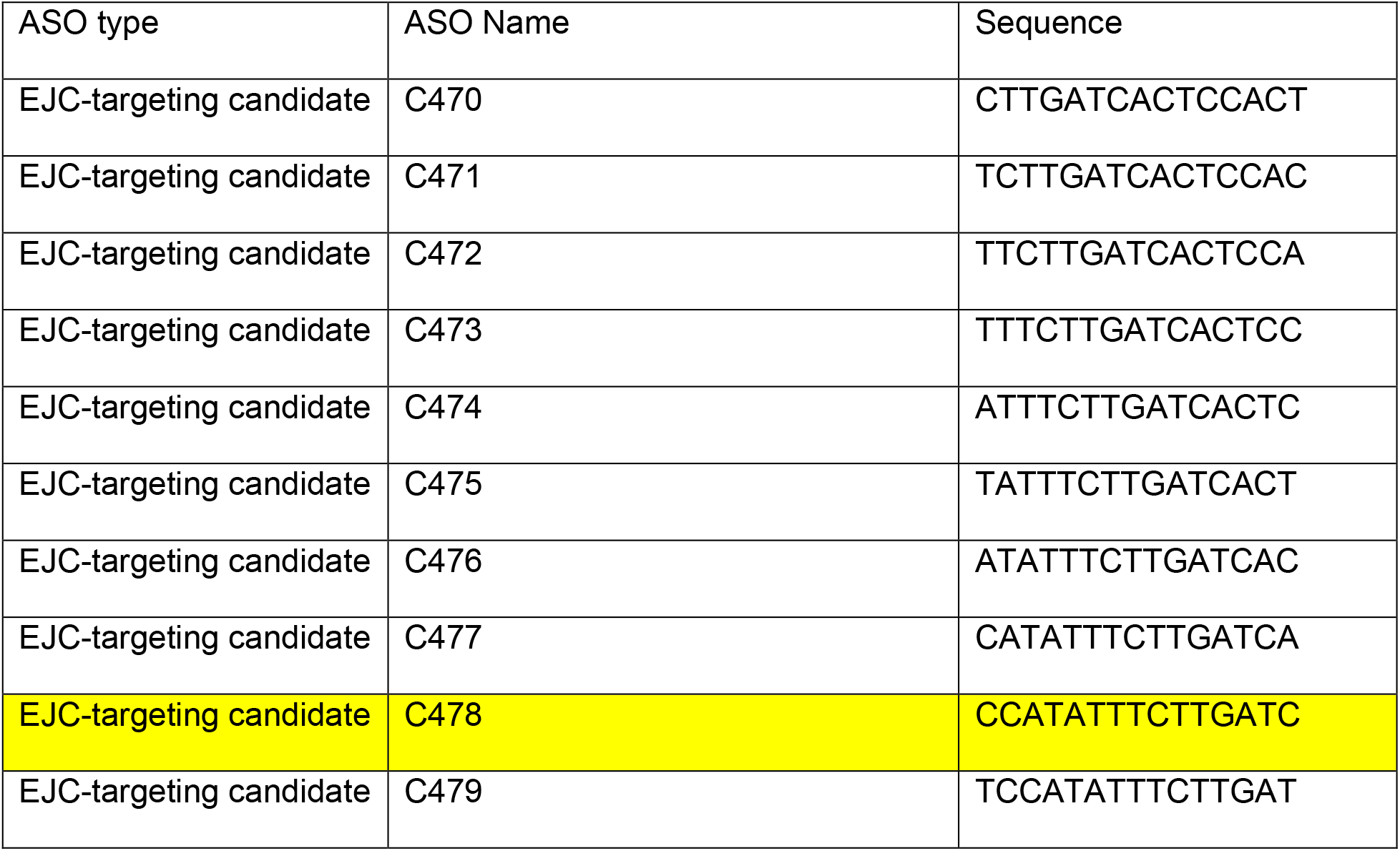
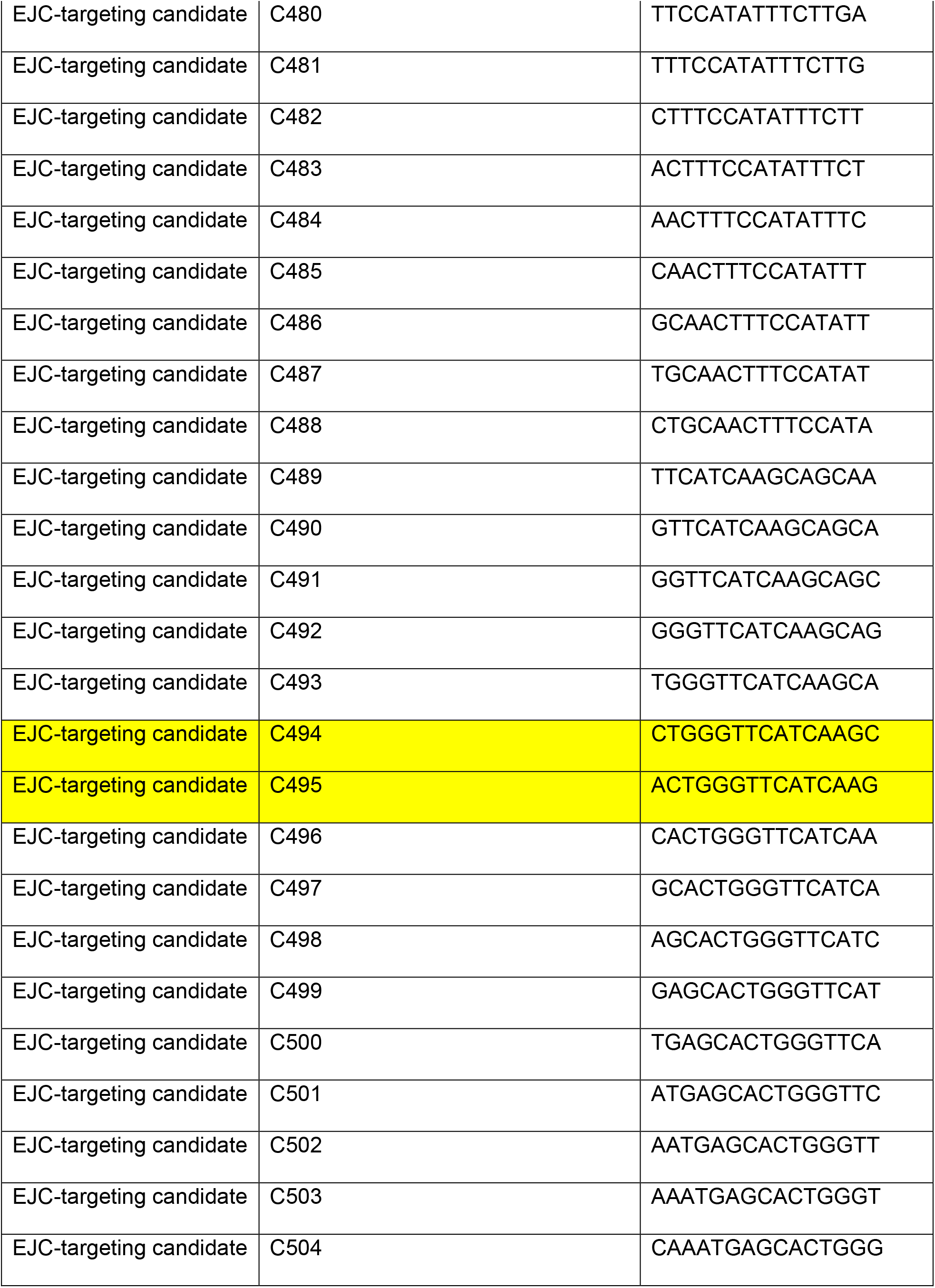
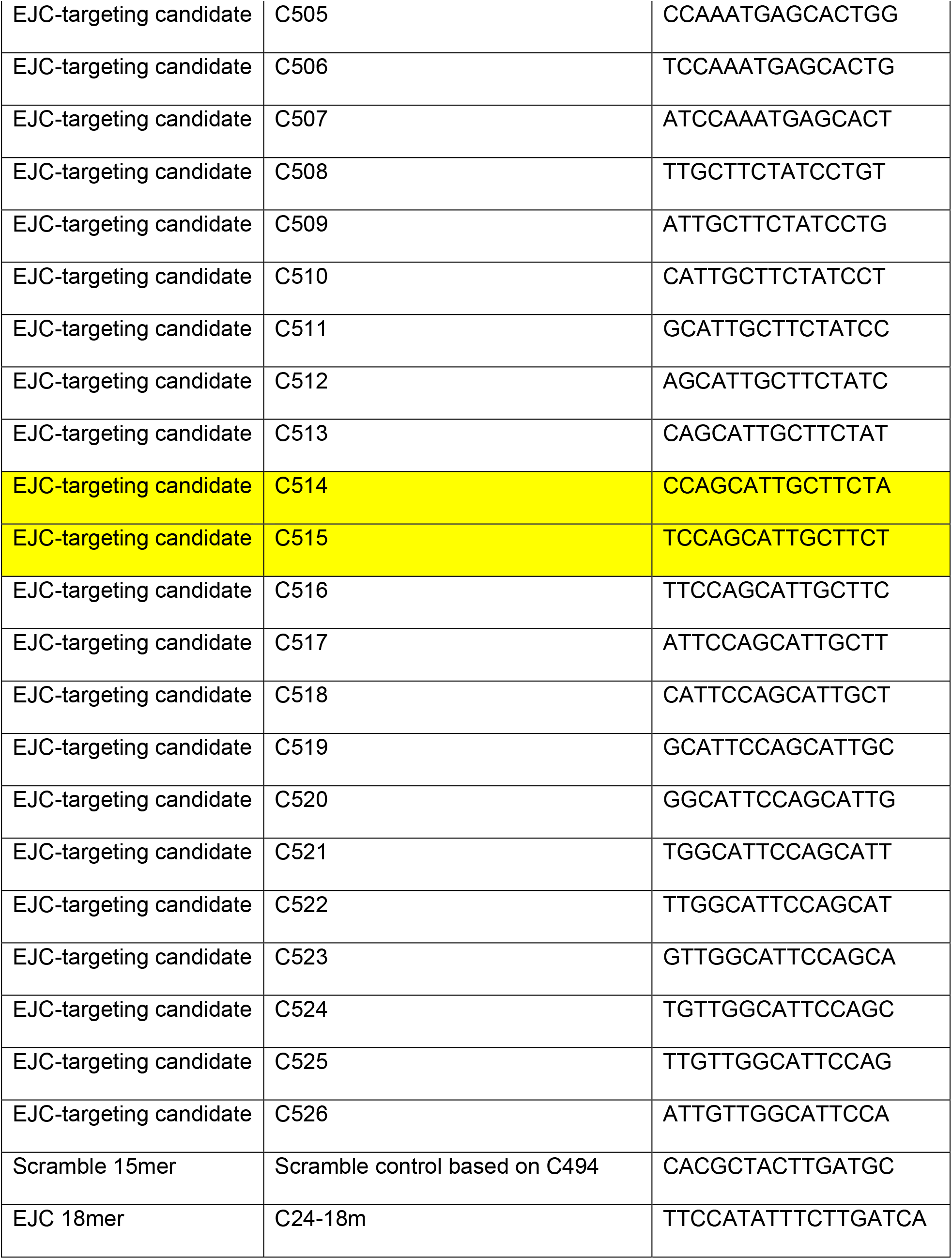
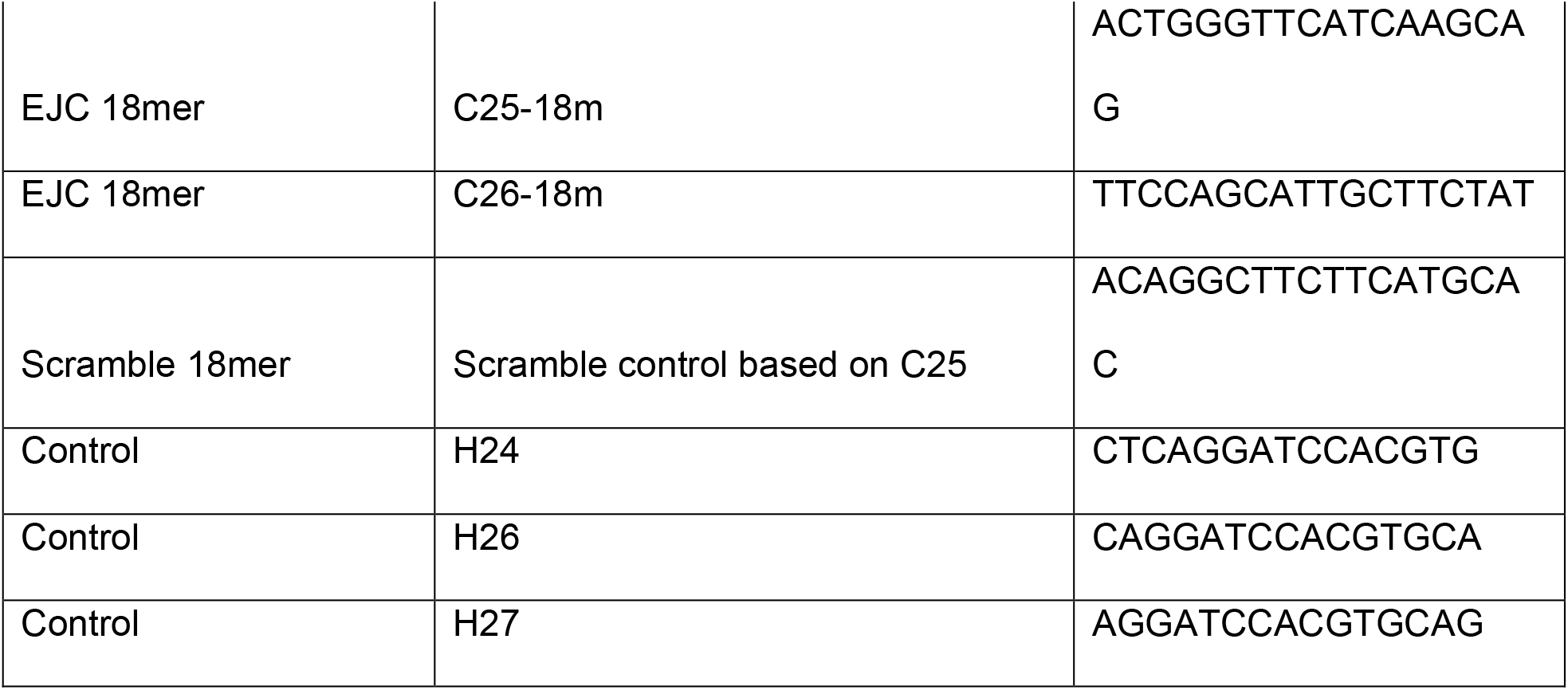
ASOs.

### Preparation of U2OS cells expressing NMD reporters

The NMD reporters used for the ASO screening (*pW1282X-IVS23, pW1282X-IVS24, pW1282X-IVS25*, and *pW1282X-IVS26*) were constructed from the parent NMD reporter *GFP-CFTR22-27-T7*, which was cloned into the pCDNA5 FRT/TO plasmid (Life Technologies, Carlsbad, CA).

pCDNA5 FRT/TO allows tetracycline-inducible expression of the gene. *GFP-CFTR22-27-T7* has the natural sequences of exons 22 to 27 of the human *CFTR* gene and shortened intervening sequences (IVS) modified from the natural sequences of introns 22 to 26 by taking 200 nucleotides (nt) from the 5’ and 3’ ends of the corresponding introns. GFP and T7 cDNA sequences were added to the 5’ and 3’ ends of each reporter, respectively, to facilitate gene, transcript, and protein detection. NMD reporters *pW1282X-IVS23, pW1282X-IVS24, pW1282X-IVS25*, and *pW1282X-IVS26* comprise only IVS23, IVS24, IVS25, or IVS26, respectively, downstream of the PTC. The 5’ss and 3’ss of IVS25 and the 3’ss of IVS23 in *pW1282X-IVS25* were mutated to stronger splice-site sequences (Supplementary Figure 2B) to promote proper splicing in the minigene context.

For stable expression of NMD reporters, the reporter plasmids were co-transfected with the pOG44 helper vector to express Flp recombinase into U2OS-TREx cells harboring a single FRT recombination site (Life Technologies). Cells with successful NMD-reporter integration were selected by hygromycin resistance. The expression and splicing of the transgenes were assessed by radioactive RT-PCR, following induction with 1 μg/ml doxycycline (Research Products International Corp, D43020-100).

### CRISPR mutant DLD1 and 16HBEge cells

Using CRISPR/Cas9, we generated DLD1 cells with homozygous *CFTR-W1282X* mutation. sgRNA against exon 23 (Table 2) was cloned downstream of the U6 promoter of the pSpCas9(BB)-2A-GFP (PX458) plasmid (Addgene plasmid # 48138) ^61^, creating pSC2G-CFTR23, which allows co-expression of the sgRNA and *Streptococcus pyogenes* Cas9 (spCas9). 4 μg of pSC2G-CFTR23 plasmid was co-transfected with 1 μM single-stranded DNA repair template (synthesized by Sigma, Table 3) comprising the *CFTR-W1282X* and silent protospacer adjacent motif (PAM) mutations, using lipofectamine 2000. Following transfection, GFP+ DLD1 cells were collected using an ARIA-I cell sorter (BD), and individual clones of cells were isolated by limiting dilution into 96-well plates. Clonal cells were expanded and passaged until confluent in 6-well plates. We characterized 159 clones by Sanger sequencing, and identified two heterozygous and two homozygous *W1282X* mutant clones. 16HBE14o-parental cells gene-edited to yield 16HBEge cell lines CFF-16HBEge CFTR W1282X, F508del, G551D, G542X, or R1162X, homozygous for *CFTR-W1282X, F508del, G551D, G542X, or R1162X* mutation, respectively, in the endogenous loci were kindly provided by the Cystic Fibrosis Foundation’s CFFT Lab ^18^. Elsewhere in the text, these cells are referred to as 16HBE-W1282X, 16HBE-F508del, 16HBE-G551D, 16HBE-G542X, or 16HBE-R1162X cells, respectively.

**Table 2.**
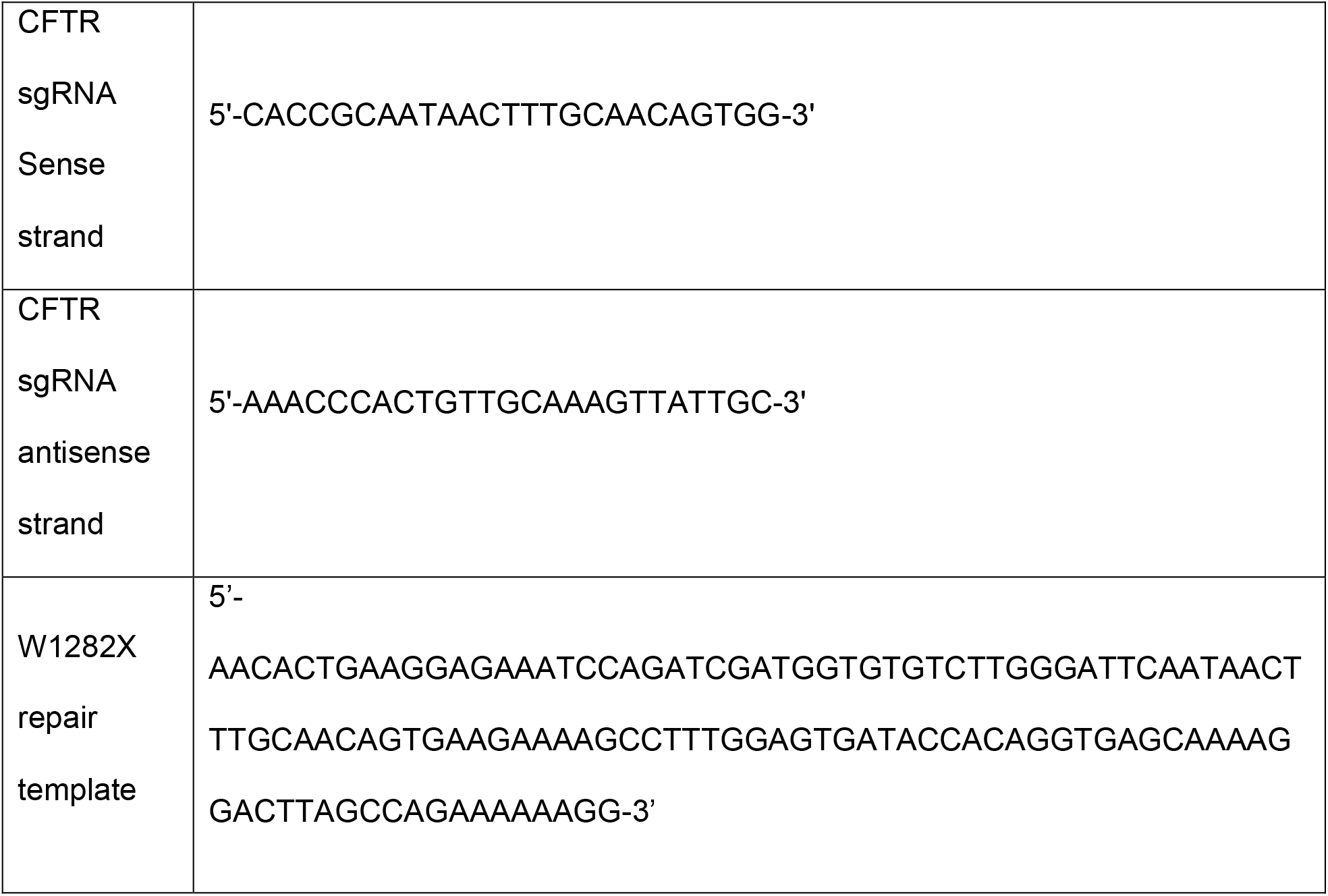
sgRNA and W1282X repair template sequences.

**Table 3.**
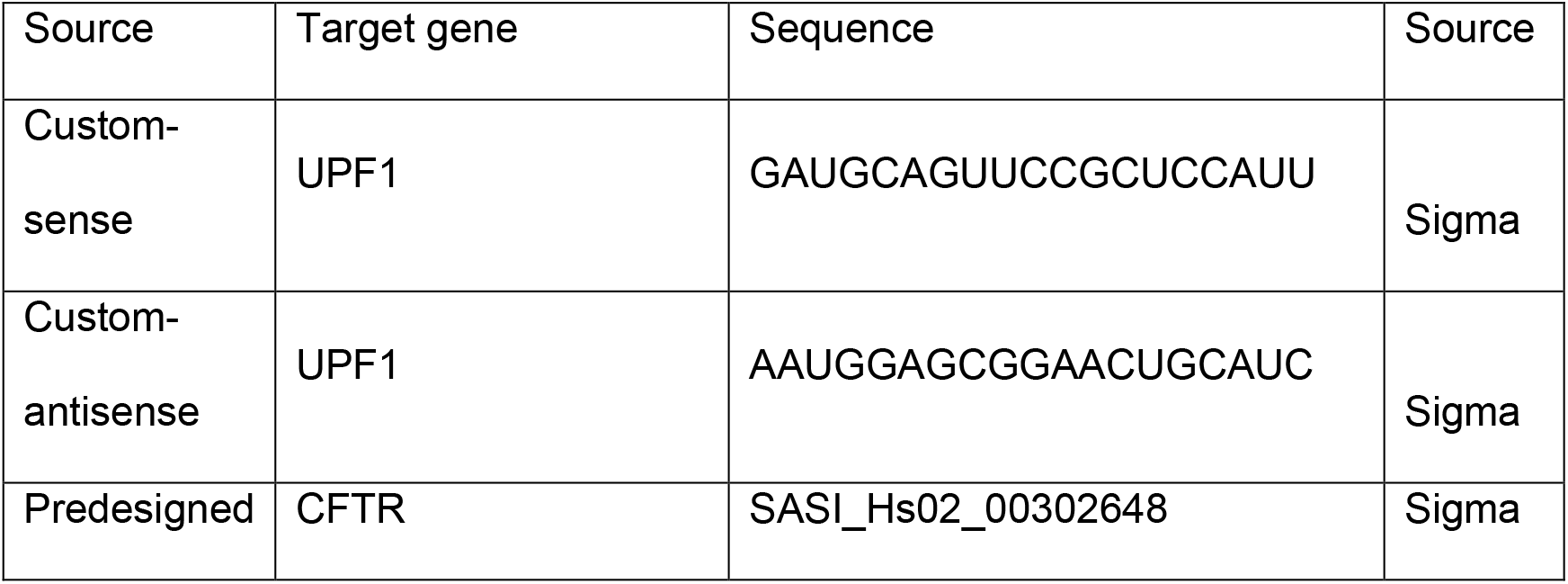
siRNA.

### Tissue culture and transfection of siRNA, ASO, and plasmids

U2OS and DLD1 cells were cultured in DMEM with 10% FBS. 16HBEge cells were cultured in MEM with 10% FBS. All cells were incubated at 37 °C and 5% CO_2_. Cells were transfected with ASOs and plasmids using Lipofectamine 3000 (Life Technologies, L3000015) according to the manufacturer’s protocol, and harvested 48 hrs post-transfection. Cells were transfected with siRNA (Table 3) using Lipofectamine RNAiMax (Life Technologies, 13778075) according to the manufacturer’s protocol for transfecting short oligonucleotides, and harvested 48 hrs post-transfection. NMD inhibition by cycloheximide was performed by treating the cells for 1 hr with cycloheximide (Sigma, 100 μg/mL). For ASO treatment by free-uptake, 1 mM stock ASO solutions were diluted into MEM with 10% FBS to the desired final concentrations, and the cells were cultured for 4 days before harvesting or Ussing-chamber assays. NMD-reporter expression was induced with 1 μg/ml doxycycline with media change, 6 hr after transfection. For the G418-treatment group, G418 (Sigma) was added to the culture medium at the indicated final concentrations, 24 hr before protein extraction or Ussing-chamber assays.

### RNA extraction and RT-PCR

Total RNA was extracted with TRIzol (Life Technologies) according to the manufacturer’s protocol. Oligo dT(18)-primed reverse transcription was carried out with ImProm-II Reverse Transcriptase (Roche). Semi-quantitative radioactive PCR (RT-PCR) was carried out in the presence of ^32^P-dCTP with AmpliTaq DNA polymerase (Thermo Fisher), and real-time quantitative RT-PCR (RT-qPCR) was performed with Power Sybr Green Master Mix (Thermo Fisher). Primers used for RT-PCR and RT-qPCR are listed in Table 4. RT-PCR products were separated by 6% native polyacrylamide gel electrophoresis, detected with a Typhoon FLA7000 phosphorimager, and quantitated using MultiGauge v2.3 software (Fujifilm); RT-qPCR data were quantitated using QuantStudio 6 Flex system.

**Table 4.**
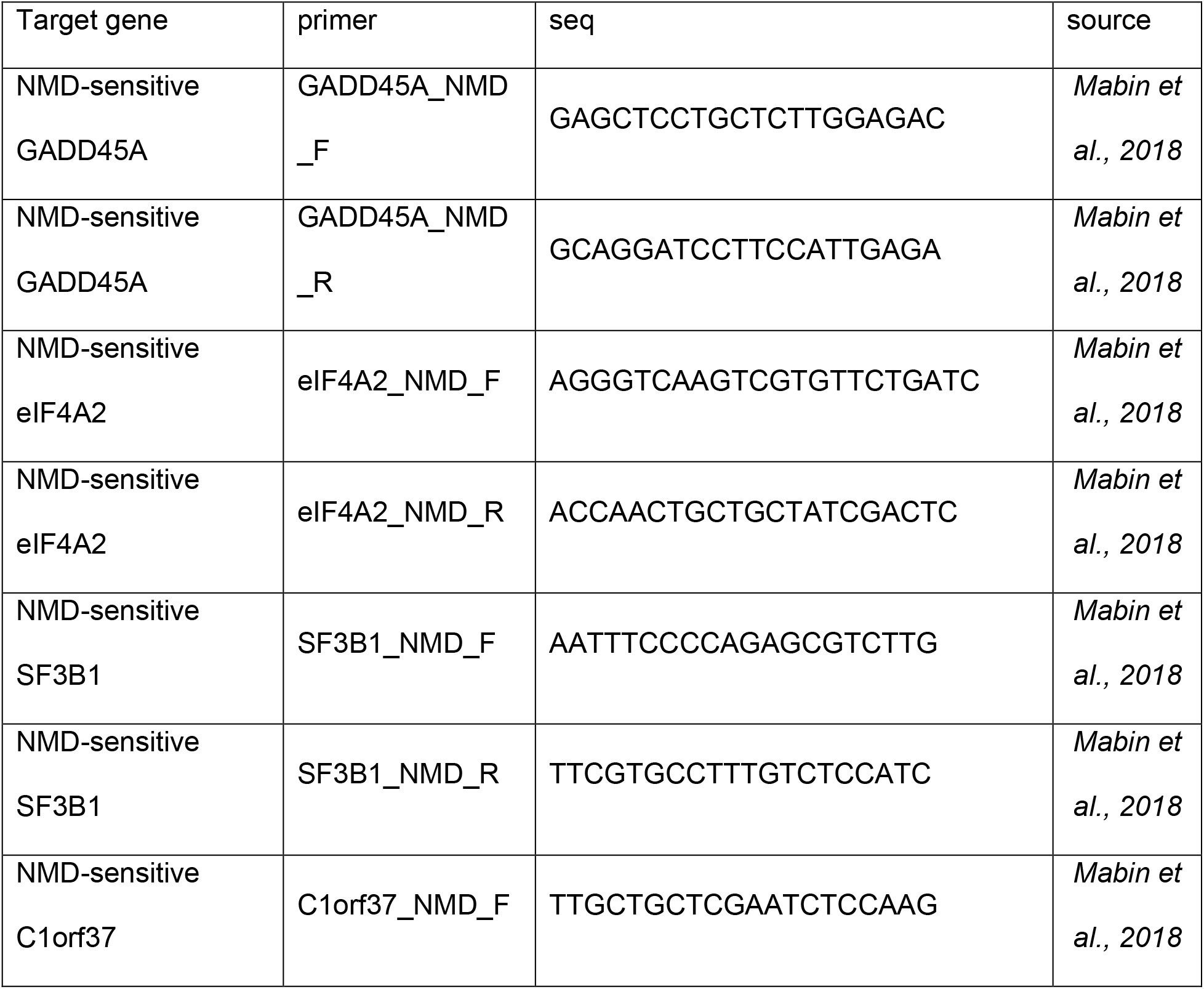
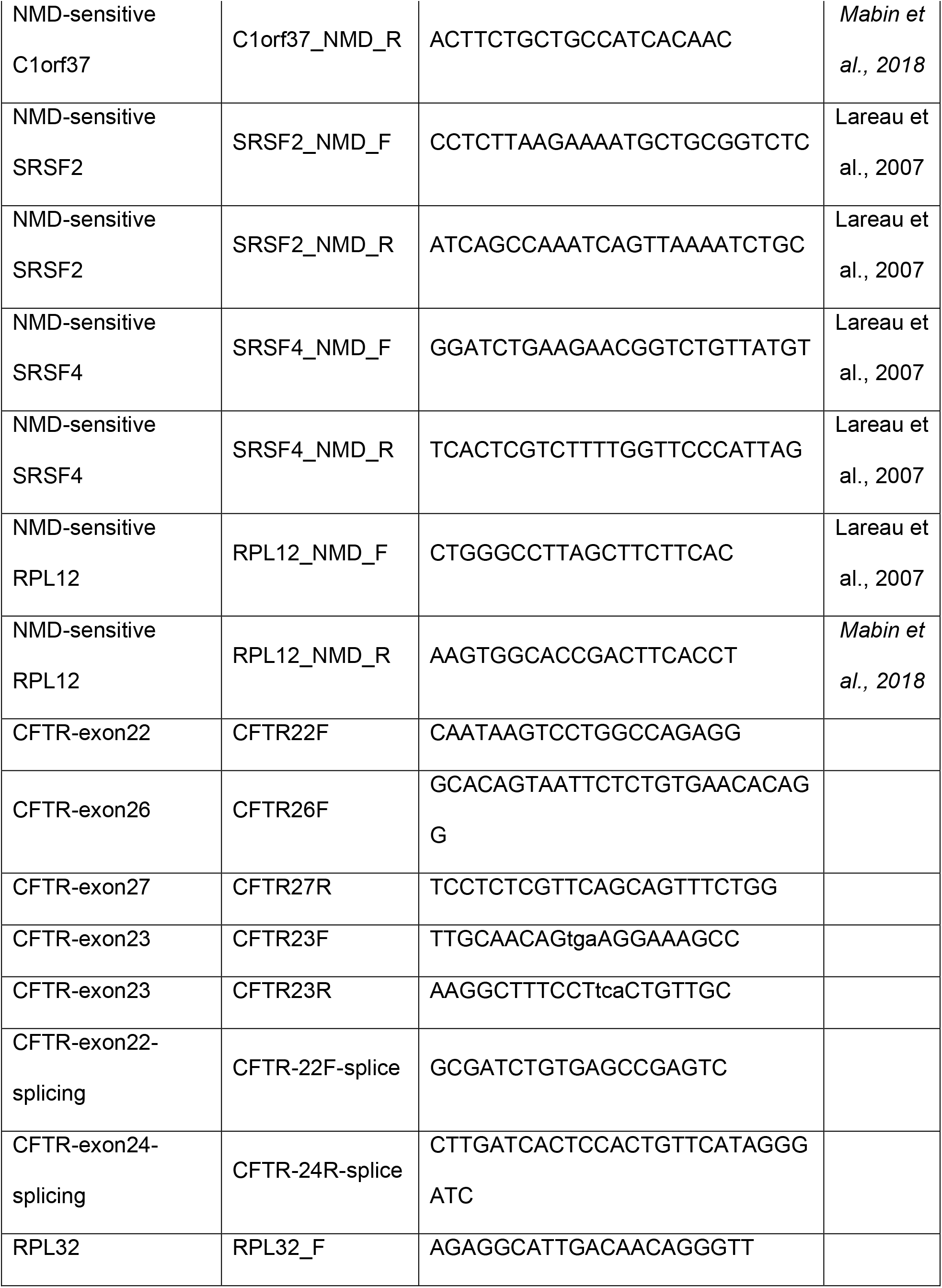
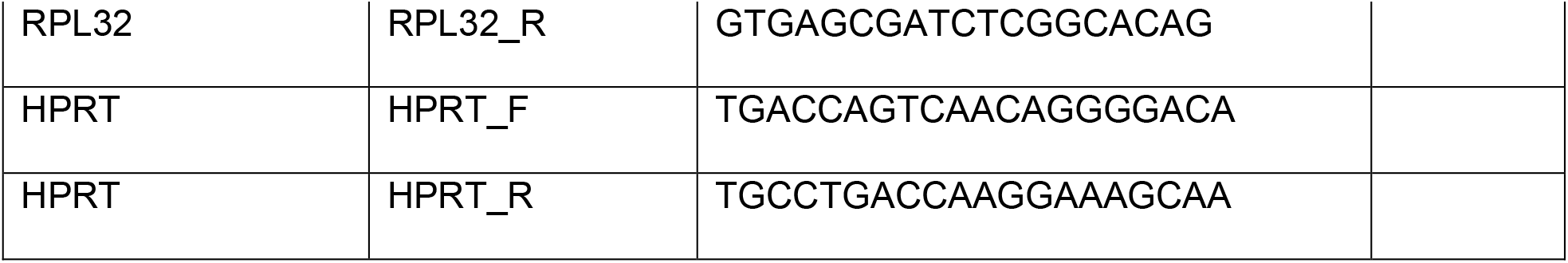
Primers for RT-PCR and RT-qPCR.

### Protein extraction, deglycosylation, and Western blotting

Cells were harvested with RIPA buffer (150 mM NaCl, 50 mM Tris-HCl pH 8.0, 1% NP40, 0.5% sodium deoxycholate, and 0.1% SDS) and 2 mM EDTA + protease inhibitor cocktail (Roche) by sonicating for 5 min at medium power using a Bioruptor (Diagenode), followed by 15-min incubation on ice. Protein concentration was measured using the Bradford assay (Bio-Rad) with BSA as a standard. To monitor post-translational maturation of CFTR protein, cell lysates were incubated in 40 μg/ml PNGase F (New England Biolabs, MA) for 2 hr at 37 °C to cleave all N-glycans before immunoblotting ^62^. Cell lysates were mixed with Laemmli buffer and incubated at 37 °C for 30 min. The protein extracts were separated by sodium dodecyl sulfate-polyacrylamide gel electrophoresis (SDS-PAGE) (6% Tris-chloride gels) and then transferred onto a nitrocellulose membrane. CFTR bands C, B, and A were detected with antibody UNC-596 (J. Riordan lab, University of North Carolina, Chapel Hill, NC). The C and B band intensities were measured together for the quantification of CFTR protein levels. The specificity of the antibody was confirmed by knocking down *CFTR* in WT DLD1 cells (Supplementary Fig. 11A). Na/K-ATPase, detected with a specific antibody (Santa Cruz sc-48345), was used as a loading control. UPF1 was detected with rabbit antibody D15G6 (Cell Signaling Technology #12040S); mouse anti-α-tubulin antibody (Sigma T9026) was used as a loading control. IRDye 800CW or 700CW secondary antibody (LI-COR) was used for Western blotting, and the blots were imaged and quantified using an Odyssey Infrared Imaging System (LI-COR). Statistical significance was calculated using Student’s *t*-test or one-way ANOVA, followed by Tukey’s or Dunnett’s post-test.

### Ussing-chamber assay

#### Preparing 16HBEge cells for Ussing-chamber assay

16HBEge cells were grown as an electrically tight monolayer on Snapwell filter supports (Corning, cat# 3801), as described ^63^, and both serosal and mucosal membranes were exposed to the ASOs for 4 days, and to CFTR correctors for 24 hrs, before the assays. The Snapwell inserts were transferred to an Ussing chamber (P2302, Physiologic Instruments, Inc., San Diego, CA). For 16HBEge cells, the serosal side only was superfused with 5mL of HB-PS buffer; on the mucosal side, 5 ml of CF-PS was used (137 mM Na-gluconate; 4 mM KCl; 1.8 mM CaCl_2_; 1 mM MgCl_2_; 10 mM HEPES; 10 mM glucose; pH adjusted to 7.4 with N-methyl-D-glucamine) to create a transepithelial chloride-ion gradient. After clamping transepithelial voltage to 0 mV, the short-circuit current (I_SC_) was measured with a Physiologic Instruments VCC MC6 epithelial voltage clamp, while maintaining the buffer temperature at 37 °C. Baseline activity was recorded for 20 min before agonists (final concentrations: 10 µM forskolin (Sigma, F6886), 50 µM genistein (Sigma, G6649), and 1-10 µM VX-770 (Selleckchem, S1144) and inhibitor (final concentration: 20 µM CFTRinh-172 (Sigma, C2992)) were applied sequentially at 10 or 20-minute intervals, to both serosal and mucosal surfaces. Agonists/inhibitor were added from 200x-1000x stock solutions. Data acquisition performed using ACQUIRE & ANALYZE Revision II (Physiologic Instruments).

### ESE motif analysis

Potential SR protein binding sites were analyzed by ESEfinder ^46^.

### Statistical analyses

Statistical analyses were performed with GraphPad Prism 5. Statistical parameters are indicated in the figures and legends. For two-tailed t-test or one-way analysis of variance (ANOVA) with Tukey’s or Dunnett’s post-test, P<0.05 was considered significant. The Pearson correlation and P values were calculated using R. The asterisks and hash signs mark statistical significance as follows: n.s. P>0.05; *^/#^ P<0.05; **^/##^ P<0.01; ***^/###^ P<0.001.

## Acknowledgments

We are very grateful to Martin Mense and Hermann Bihler (CFFT, Lexington, MA) for generously sharing protocols and advice, and for helpful comments on the manuscript.

## Author contribution

Y.J.K., T.N., and A.R.K. conceived the study. A.R.K. supervised the study. Y.J.K. and T.N. generated the DLD1-W1282X cells. F.P. performed exon 25-targeting ASO screening using the *pW1282X-IVS25* NMD reporter. Y.J.K. designed and performed all other experiments and analyzed the data. Y.J.K. and A.R.K. wrote the paper, and all authors approved the manuscript.

## References

1. CFTR2. Clinical and Functional Translation of CFTR (CFTR2). https://cftr2.org/ (2020).

2. Cystic Fibrosis Foundation. Patient Registry Annual Data Report. Lung http://www.cff.org/UploadedFiles/research/ClinicalResearch/Patient-Registry-Report-2009.pdf (2018).

3. Abeliovich, D. et al. Screening for five mutations detects 97% of cystic fibrosis (CF) chromosomes and predicts a carrier frequency of 1:29 in the Jewish Ashkenazi population. Am. J. Hum. Genet. 51, 951–6 (1992).

4. Rowe, S. M. et al. Restoration of W1282X CFTR Activity by Enhanced Expression. Am. J. Respir. Cell Mol. Biol. 37, 347–356 (2007).

5. Haggie, P. M. et al. Correctors and potentiators rescue function of the truncated W1282X-CFTR translation product. J. Biol. Chem. 292, jbc.M116.764720 (2016).

6. Wang, W., Hong, J. S., Rab, A., Sorscher, E. J. & Kirk, K. L. Robust stimulation of W1282X-CFTR channel activity by a combination of allosteric modulators. PLoS One 11, e0152232 (2016).

7. Goor, F. Van et al. Rescue of CF airway epithelial cell function in vitro by a CFTR potentiator, VX-770. Proc Natl Acad Sci U S A 106, 18825–18830 (2009).

8. Johnson, L. G. et al. Efficiency of gene transfer for restoration of normal airway epithelial function in cystic fibrosis. Nat. Genet. 2, 21–25 (1992).

9. Keating, D. et al. VX-445-Tezacaftor-Ivacaftor in patients with cystic fibrosis and one or two Phe508del alleles. N. Engl. J. Med. 379, 1612–1620 (2018).

10. Lentini, L. et al. Toward a rationale for the PTC124 (Ataluren) promoted readthrough of premature stop codons: a computational approach and GFP-reporter cell-based assay. Mol. Pharm. 11, 653–664 (2014).

11. Linde, L. et al. Nonsense-mediated mRNA decay affects nonsense transcript levels and governs response of cystic fibrosis patients to gentamicin. J. Clin. Invest. 117, 683–692 (2007).

12. Clancy, J. P. et al. No detectable improvements in cystic fibrosis transmembrane conductance regulator by nasal aminoglycosides in patients with cystic fibrosis with stop mutations. Am. J. Respir. Cell Mol. Biol. 37, 57–66 (2007).

13. Kerem, E. et al. Ataluren for the treatment of nonsense-mutation cystic fibrosis: A randomised, double-blind, placebo-controlled phase 3 trial. Lancet Respir. Med. 2, 539– 547 (2014).

14. Zainal Abidin, N., Haq, I. J., Gardner, A. I. & Brodlie, M. Ataluren in cystic fibrosis: development, clinical studies and where are we now? Expert Opin. Pharmacother. 18, 1363–1371 (2017).

15. Keeling, K. M. et al. Attenuation of nonsense-mediated mRNA decay enhances in vivo nonsense suppression. PLoS One 8, e60478 (2013).

16. Mutyam, V. et al. Therapeutic benefit observed with the CFTR potentiator, ivacaftor, in a CF patient homozygous for the W1282X CFTR nonsense mutation. J. Cyst. Fibros. (2016) doi:10.1016/j.jcf.2016.09.005.

17. Usuki, F. et al. Inhibition of SMG-8, a subunit of SMG-1 kinase, ameliorates nonsense-mediated mRNA decay-exacerbated mutant phenotypes without cytotoxicity. Proc. Natl. Acad. Sci. 110, 15037–15042 (2013).

18. Valley, H. C. et al. Isogenic cell models of cystic fibrosis-causing variants in natively expressing pulmonary epithelial cells. J. Cyst. Fibros. 8–15 (2018) doi:10.1016/j.jcf.2018.12.001.

19. Keenan, M. M. et al. Nonsense-mediated RNA decay pathway inhibition restores expression and function of W1282X CFTR. Am. J. Respir. Cell Mol. Biol. 61, 290–300 (2019).

20. Huang, L. et al. Antisense suppression of the nonsense mediated decay factor Upf3b as a potential treatment for diseases caused by nonsense mutations. Genome Biol. 19, 1–16 (2018).

21. Lykke-Andersen, S. & Jensen, T. H. Nonsense-mediated mRNA decay: an intricate machinery that shapes transcriptomes. Nat. Rev. Mol. Cell Biol. (2015) doi:10.1038/nrm4063.

22. Singh, G. et al. The cellular EJC interactome reveals higher order mRNP structure and an EJC-SR protein nexus. Cell 151, 750–764 (2012).

23. Le Hir, H., Saulière, J. & Wang, Z. The exon junction complex as a node of post-transcriptional networks. Nat. Rev. Mol. Cell Biol. 17, 41–54 (2016).

24. Haque, N. & Blanchette, M. Genomewide localization of Exon-Junction-Complex (EJC) in Drosophila. FASEB J. 25, 510.1-510.1.

25. Saulière, J. et al. CLIP-seq of eIF4AIII reveals transcriptome-wide mapping of the human exon junction complex. Nat. Struct. Mol. Biol. 19, 1124–1131 (2012).

26. Hauer, C. et al. Exon Junction Complexes Show a Distributional Bias toward Alternatively Spliced mRNAs and against mRNAs Coding for Ribosomal Proteins Article Exon Junction omplexes Show a Distributional Bias toward Alternatively Spliced mRNAs and against mRNAs Coding fo. CellReports 1–16 (2016) doi:10.1016/j.celrep.2016.06.096.

27. Brogna, S. & Wen, J. Nonsense-mediated mRNA decay (NMD) mechanisms. Nat. Struct. Mol. Biol. 16, 107–113 (2009).

28. Hug, N., Longman, D. & Cáceres, J. F. Mechanism and regulation of the nonsense-mediated decay pathway. Nucleic Acids Res. 44, 1483–1495 (2016).

29. Nomakuchi, T. T., Rigo, F., Aznarez, I. & Krainer, A. R. Antisense oligonucleotide-directed inhibition of nonsense-mediated mRNA decay. Nat Biotech 34, 164–166 (2016).

30. Gogtay, N. J. & Sridharan, K. Therapeutic Nucleic Acids: Current clinical status. Br. J. Clin. Pharmacol. 1–14 (2016) doi:10.1111/bcp.12987.

31. Khvorova, A. & Watts, J. K. The chemical evolution of oligonucleotide therapies of clinical utility. Nat. Biotechnol. 35, 238–248 (2017).

32. Lundin, K. E., Gissberg, O. & Smith, C. I. E. Oligonucleotide therapies: the past and the present. Hum. Gene Ther. 26, 475–485 (2015).

33. Khoo, B., Roca, X., Chew, S. L. & Krainer, A. R. Antisense oligonucleotide-induced alternative splicing of the APOB mRNA generates a novel isoform of APOB. BMC Mol. Biol. 8, 3 (2007).

34. Kole, R., Krainer, A. R. & Altman, S. RNA therapeutics: beyond RNA interference and antisense oligonucleotides. Nat. Rev. Drug Discov. 11, 125–140 (2012).

35. Shimo, T., Maruyama, R. & Yokota, T. Designing effective antisense oligonucleotides for exon skipping. Methods Mol. Biol. 1687, 143–155 (2018).

36. Shimo, T. et al. Design and evaluation of locked nucleic acid-based splice-switching oligonucleotides in vitro. Nucleic Acids Res. 42, 8174–8187 (2014).

37. Harding, P. L., Fall, A. M., Honeyman, K., Fletcher, S. & Wilton, S. D. The influence of antisense oligonucleotide length on dystrophin exon skipping. Mol. Ther. 15, 157–166 (2007).

38. Mabin, J. W. et al. The exon junction complex undergoes a compositional switch that alters mRNP structure and nonsense-mediated mRNA decay activity. Cell Rep. 25, 1–16 (2018).

39. Lareau, L. F., Inada, M., Green, R. E., Wengrod, J. C. & Brenner, S. E. Unproductive splicing of SR genes associated with highly conserved and ultraconserved DNA elements. Nature 446, 926–929 (2007).

40. Popp, M.W.-L. & Maquat, L. E. The dharma of nonsense-mediated mRNA decay in mammalian cells. Mol. Cells 37, 1–8 (2014).

41. Lima, W. F., Vickers, T. A., Nichols, J., Li, C. & Crooke, S. T. Defining the factors that contribute to on-target specificity of antisense oligonucleotides. PLoS One 9, e101752 (2014).

42. Sahashi, K. et al. TSUNAMI: An antisense method to phenocopy splicing-associated diseases in animals. Genes Dev. 26, 1874–1884 (2012).

43. Busch, A. & Hertel, K. J. Evolution of SR protein and hnRNP splicing regulatory factors. Wiley Interdiscip. Rev. RNA 3, 1–12 (2012).

44. Sinha, R. et al. Antisense oligonucleotides correct the familial dysautonomia splicing defect in IKBKAPtransgenic mice. Nucleic Acids Res. 46, 4833–4844 (2018).

45. Hua, Y., Vickers, T. A., Baker, B. F., Bennett, C. F. & Krainer, A. R. Enhancement of SMN2 Exon 7 Inclusion by Antisense Oligonucleotides Targeting the Exon. PLoS Biol. 5, e73 (2007).

46. Cartegni, L., Wang, J., Zhu, Z., Zhang, M. Q. & Krainer, A. R. ESEfinder: A web resource to identify exonic splicing enhancers. Nucleic Acids Res. 31, 3568–3571 (2003).

47. Yoshida, T. et al. Evaluation of off-target effects of gapmer antisense oligonucleotides using human cells. Genes to Cells 24, 827–835 (2019).

48. Roignant, J. Y. & Treisman, J. E. Exon junction complex subunits are required to splice Drosophila MAP kinase, a large heterochromatic gene. Cell 143, 238–250 (2010).

49. Wang, Z. et al. Transcriptome-wide modulation of splicing by the exon junction complex. Genome Biol. 15, 551 (2014).

50. Blazquez, L. et al. Exon junction complex shapes the transcriptome by repressing recursive splicing. Mol. Cell 72, 496-509.e9 (2018).

51. Boehm, V. et al. Exon junction complexes suppress spurious splice sites to safeguard transcriptome integrity. Mol. Cell 72, 482-495.e7 (2018).

52. Michelle, L. et al. Proteins associated with the exon junction complex also control the alternative splicing of apoptotic regulators. Mol. Cell. Biol. 32, 954–967 (2012).

53. Fukumura, K. et al. The exon junction complex controls the efficient and faithful splicing of a subset of transcripts involved in mitotic cell-cycle progression. Int. J. Mol. Sci. 17, (2016).

54. Saulière, J. et al. The exon junction complex differentially marks spliced junctions. Nat. Struct. Mol. Biol. 17, 1269–1271 (2010).

55. Ren, H. Y. et al. VX-809 corrects folding defects in cystic fibrosis transmembrane conductance regulator protein through action on membrane-spanning domain 1. Mol. Biol. Cell 24, 3016–3024 (2013).

56. O’Riordan, C. R., Lachapelle, A. L., Marshall, J., Higgins, E. A. & Cheng, S. H. Characterization of the oligosaccharide structures associated with the cystic fibrosis transmembrane conductance regulator. Glycobiology 10, 1225–1233 (2000).

57. Cozens, A. L. et al. CFTR expression and chloride secretion in polarized immortal human bronchial epithelial cells. Am. J. Respir. Cell Mol. Biol. 10, 38–47 (1994).

58. Xue, X. et al. Identification of the amino acids inserted during suppression of CFTR nonsense mutations and determination of their functional consequences. Hum. Mol. Genet. 26, 3116–3129 (2017).

59. Mutyam, V. et al. Discovery of clinically approved agents that promote suppression of cystic fibrosis transmembrane conductance regulator nonsense mutations. Am. J. Respir. Crit. Care Med. 194, 1092–1103 (2016).

60. Scharner, J. et al. Hybridization-mediated off-target effects of splice-switching antisense oligonucleotides. Nucleic Acids Res. 1–15 (2019) doi:10.1093/nar/gkz1132.

61. Ran, F. A. et al. Genome engineering using the CRISPR-Cas9 system. Nat. Protoc. 8, 2281–2308 (2013).

62. Glozman, R. et al. N-glycans are direct determinants of CFTR folding and stability in secretory and endocytic membrane traffic. J. Cell Biol. 184, 847–862 (2009).

63. Zegarra-Moran, O. et al. Correction of G551D-CFTR transport defect in epithelial monolayers by genistein but not by CPX or MPB-07. Br. J. Pharmacol. 137, 504–512 (2002).

