## Supplementary Figures for "Gene-Specific Nonsense-Mediated mRNA Decay Targeting for Cystic Fibrosis Therapy"

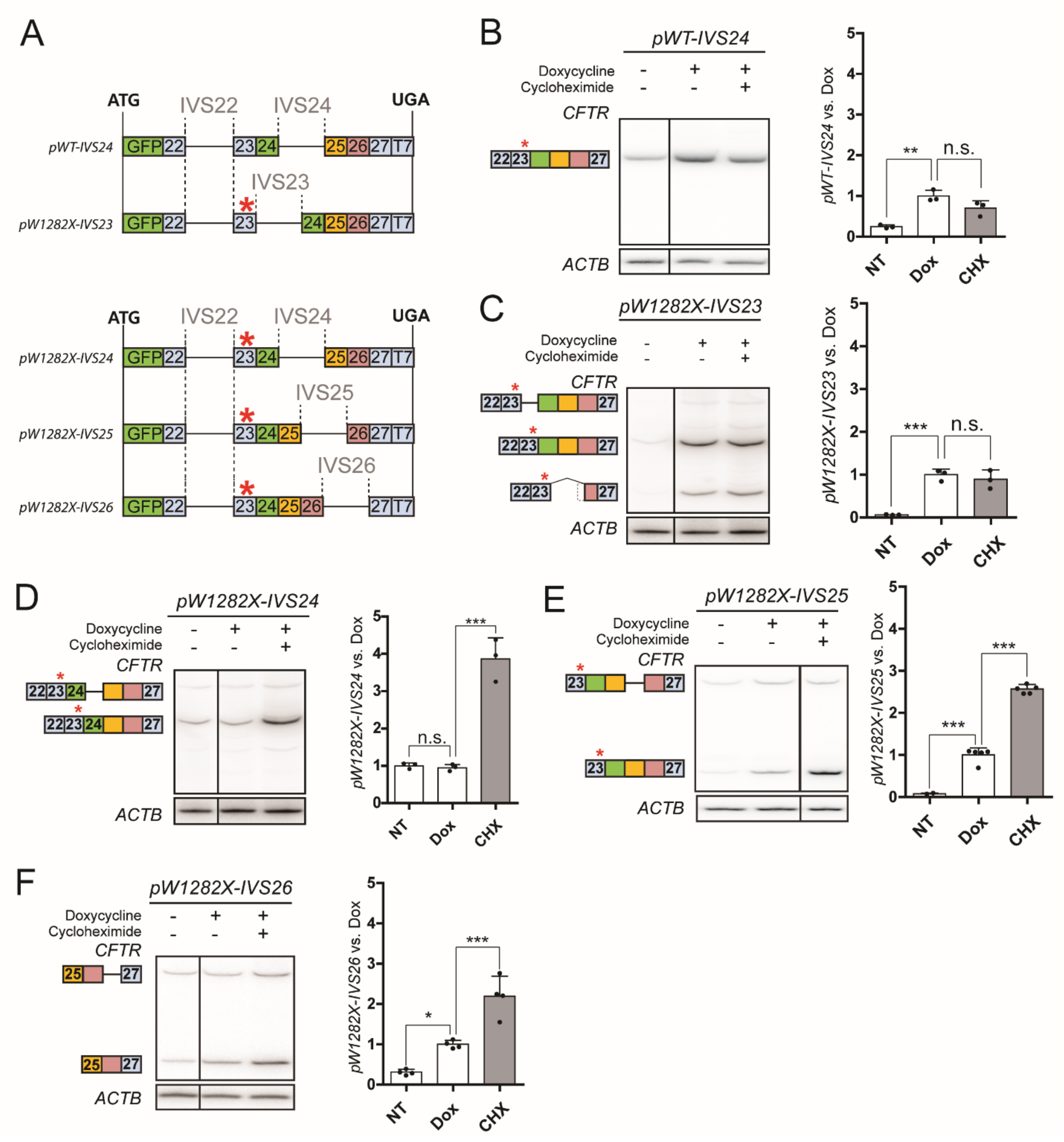


**Supplementary Figure 1. Characterization of *CFTR* NMD reporter**

**A.** Schematic of NMD reporters. The numbers in the boxes show the *CFTR* exons present in the NMD reporters. The red asterisk (*) indicates the location of the W1282X mutation. **B-F.** The radioactively labeled RT-PCR images show representative mRNA levels of NMD reporters (B) *pWT-IVS24,* (C) *pW1282X-IVS23,* (D) *pW1282X-IVS24,* (E) *pW1282X-IVS25,* and (F) *pW1282X-*

*IVS26*. *ACTB* mRNA served as internal reference. The reporter mRNA levels normalized to Dox are shown on the graph on the right side in each panel (n=3 independent measurements in panel B-D, n=2 or 5 in panel E, n=4 in panel F; n.s. $P>0.05$, *$P<0.05$, **$P<0.01$, ***$P<0.001$, one-way ANOVA with Tukey's post-test). NT: No-treatment; Dox: doxycycline 1 μg/mL; CHX: 100 μg/mL cycloheximide for 1 hr.

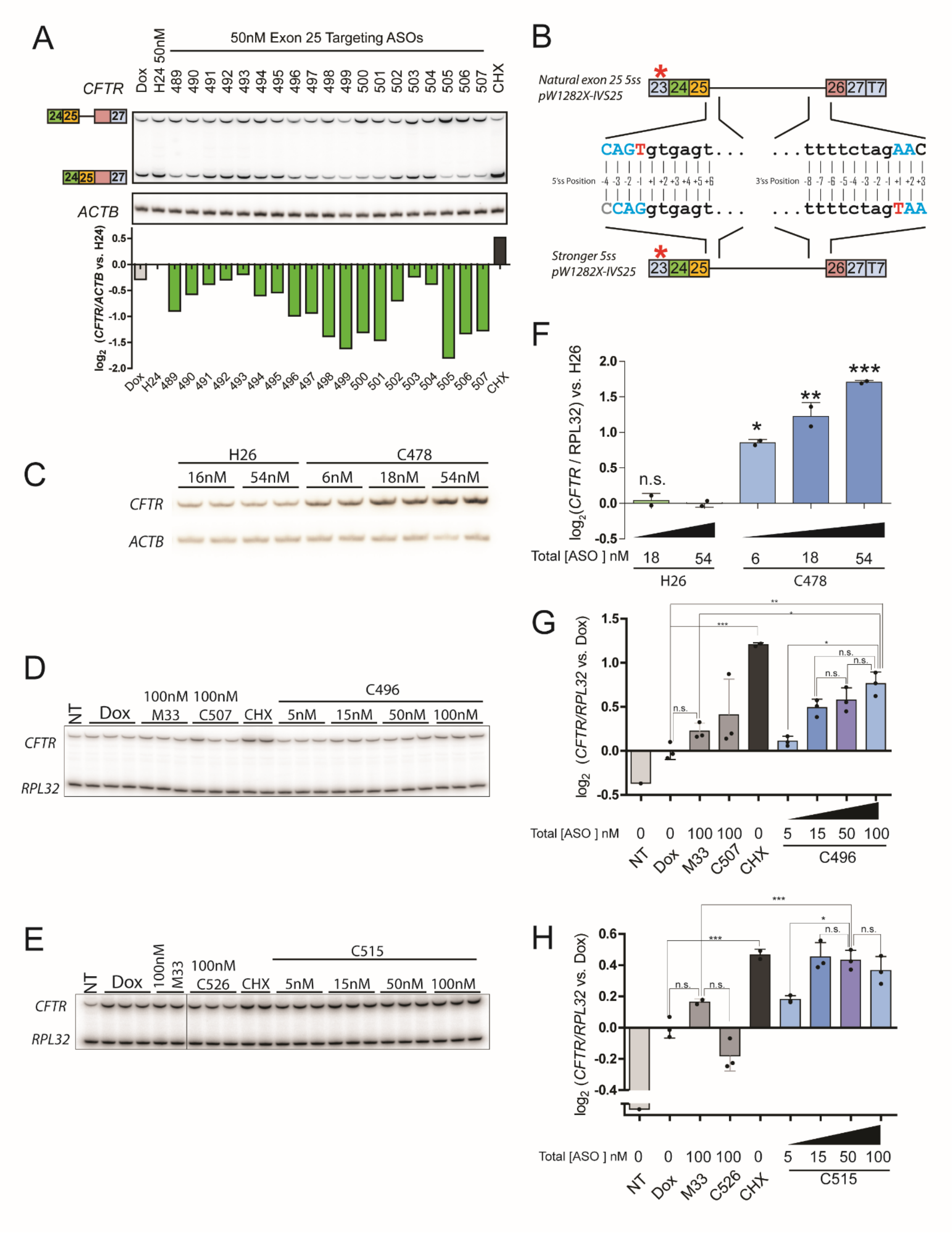


**Supplementary Figure 2. ASO screen with a weak splice-site reporter, and dose-**

**dependent increase in NMD reporter mRNAs upon candidate-ASO transfection.**

**A.** U2OS cells stably expressing *pW1282X-IVS25* with weak splice sites were transfected with 50 nM negative-control ASOs (H24) or ASOs targeting EJC-binding regions on *CFTR* exon 25, and the log2 fold changes of the reporter levels relative to the control ASO were measured by RT-PCR. *ACTB* served as internal reference. **B.** The 5'ss of *IVS25* was strengthened by deleting the T at -1, and a T was inserted at +1 of the 3'ss to preserve the same encoded protein sequence *in pW1282X-IVS25.* Upper case = exon sequence; lower case = intron sequence. **C-E.** The lead ASO candidates targeting *CFTR* (C) exon 24, (D) exon 25, or (E) exon 26 or control ASOs (H26 or M33) were tested by transfection into U2OS cells stably expressing the NMD reporters *pW1282X-IVS24, pW1282X-IVS25,* or *pW1282X-IVS26,* respectively. **F-H.** The log2 fold changes in NMD reporters in (C-E) were normalized to negative-control ASO transfection. The reporter levels were quantified by RT-PCR. *RPL32* or *ACTB* levels served as internal controls. (n=2 independent transfections for C478 and n=3 independent transfection for C496 and C515; #P < 0.05, ##P < 0.01, ###P<0.001, versus Dox, Student's t-test; *P<0.05, **P<0.01, Student's t-test). Error bars are standard deviations. Abbreviations are as in Supplemental Figure 1. H24, H26, M33: negative-control ASOs.

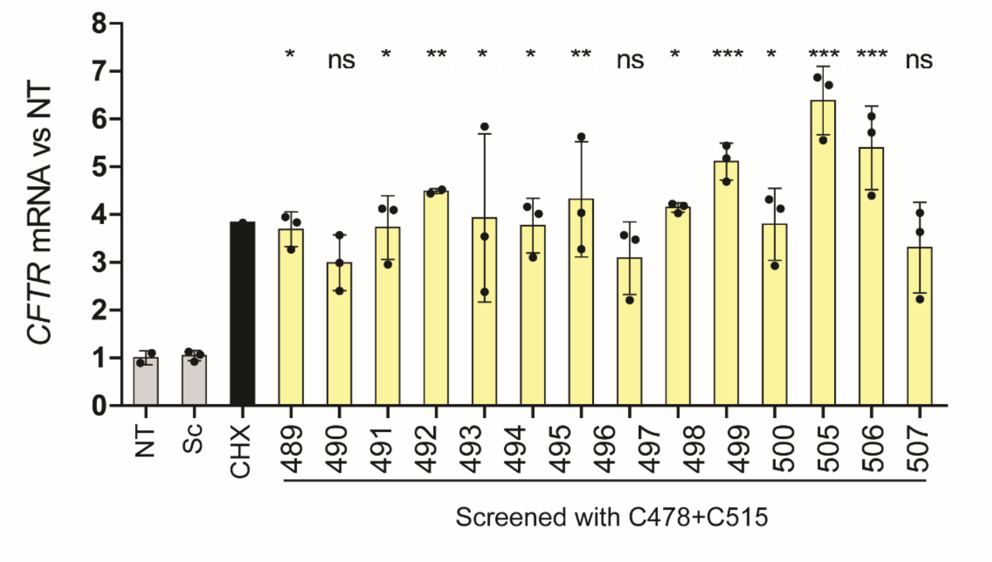


**Supplementary Figure 3. Effect of the ASO cocktail on DLD1-W1282X cells.**

Various exon-25-targeting ASOs were combined with C478 and C515, and transfected into DLD1-W1282X cells. The identity of the ASOs and nominal concentrations used for transfection are indicated. (n=3 independent treatments and transfections, n.s. $P>0.05$, *$P<0.05$, **$P<0.01$, ***$P<0.001$ versus NT, one-way ANOVA with Dunnett's post-test). All mRNA levels were measured by RT-qPCR. *RPL32* mRNA served as internal reference. Error bars show standard deviations.

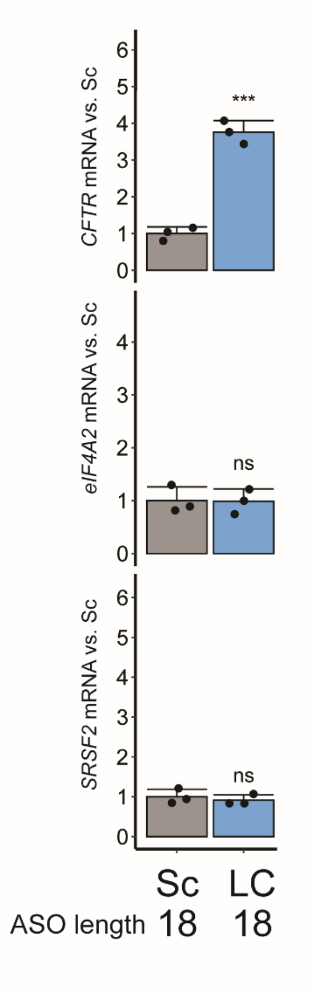


**Supplementary Figure 4. Gene-specific NMD inhibition by the 18mer ASO cocktail.**

Endogenous NMD-sensitive mRNA levels in 16HBE-W1282X cells treated with 120 nM scramble ASO (Sc) or 120 nM lead ASO cocktail C24-C25-C26 (LC) (n=3 independent treatments, n.s. P>0.05, ***P<0.001 versus NT, Student's t-test). mRNA levels were quantified by RT-qPCR. *RPL32* served as internal control. Error bars show standard deviation. Sc=Scramble ASO; LC=Lead ASO cocktail C24-C25-C26.

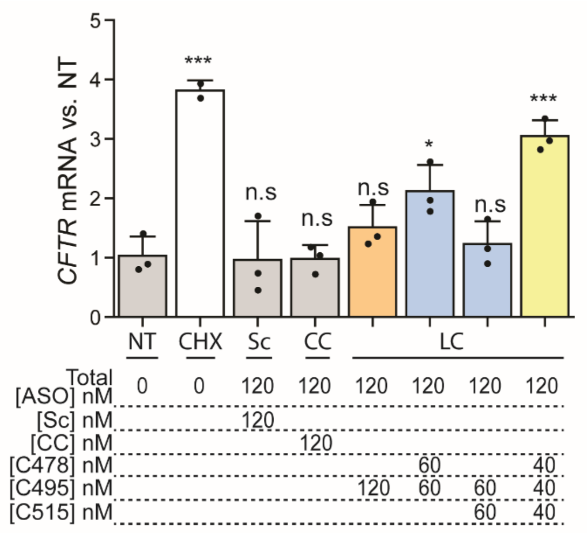


**Supplementary Figure 5. Effect of targeting different numbers of EJCs on NMD of *CFTR-W1282X* mRNA in DLD1-W1282X cells.**

The number of required EJCs targeted by the lead ASOs (C478, C495, and C515) was assessed by transfecting DLD1-W1282X cells with one, two, or three EJC-targeting ASOs at the same total nominal concentration.

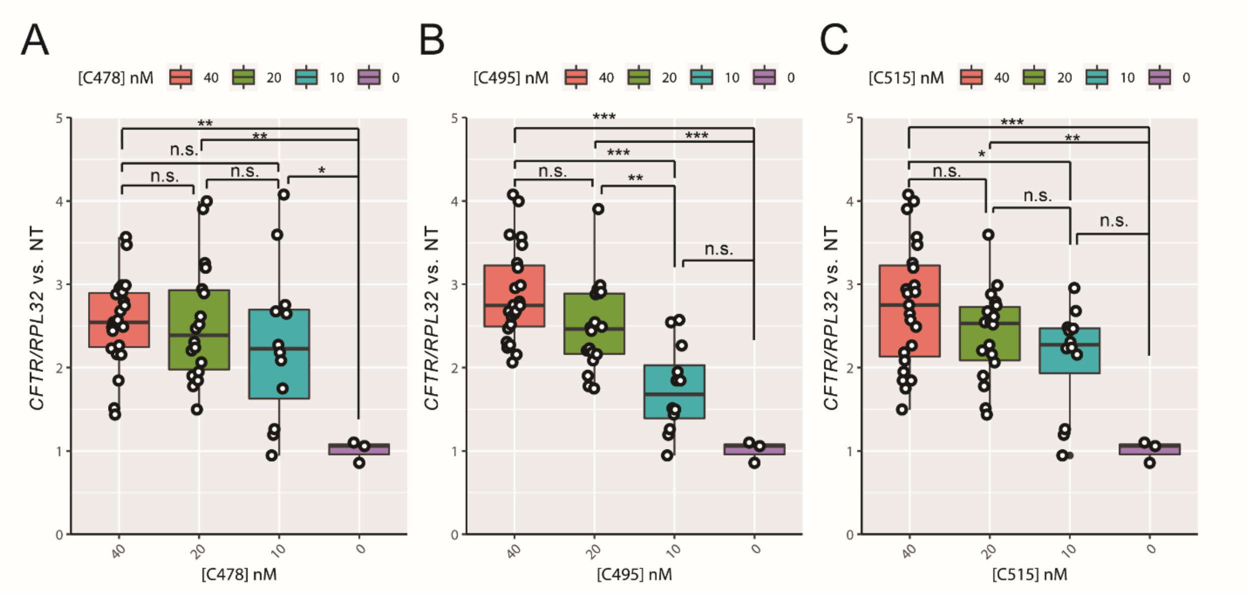


**Supplementary Figure 6. Effect of limiting ASO concentration on NMD inhibition.**

**A-C** The effect of limiting the dose of each ASO on *CFTR* mRNA levels in 16HBE-W1282X cells was quantified by categorizing the three-ASO combinations tested in Figure 2G, according to the variable concentration of **(A)** C478, **(B)** C495, and **(C)** C515 ASO. The distribution of *CFTR* mRNA levels for each ASO concentration is shown as boxplots (Box = first to third quartile; horizontal line through the box = median; Vertical lines below and above the boxes = minimum to first quartile and third quartile to maximum, respectively; individual data points plotted as open circles). Each dot represents an independent transfection of an ASO cocktail. All mRNA levels were measured by RT-qPCR. *RPL32* mRNA served as internal reference. (n.s. P>0.05, *P<0.05, **P<0.01, ***P<0.001, one-way ANOVA with Tukey's post-test).

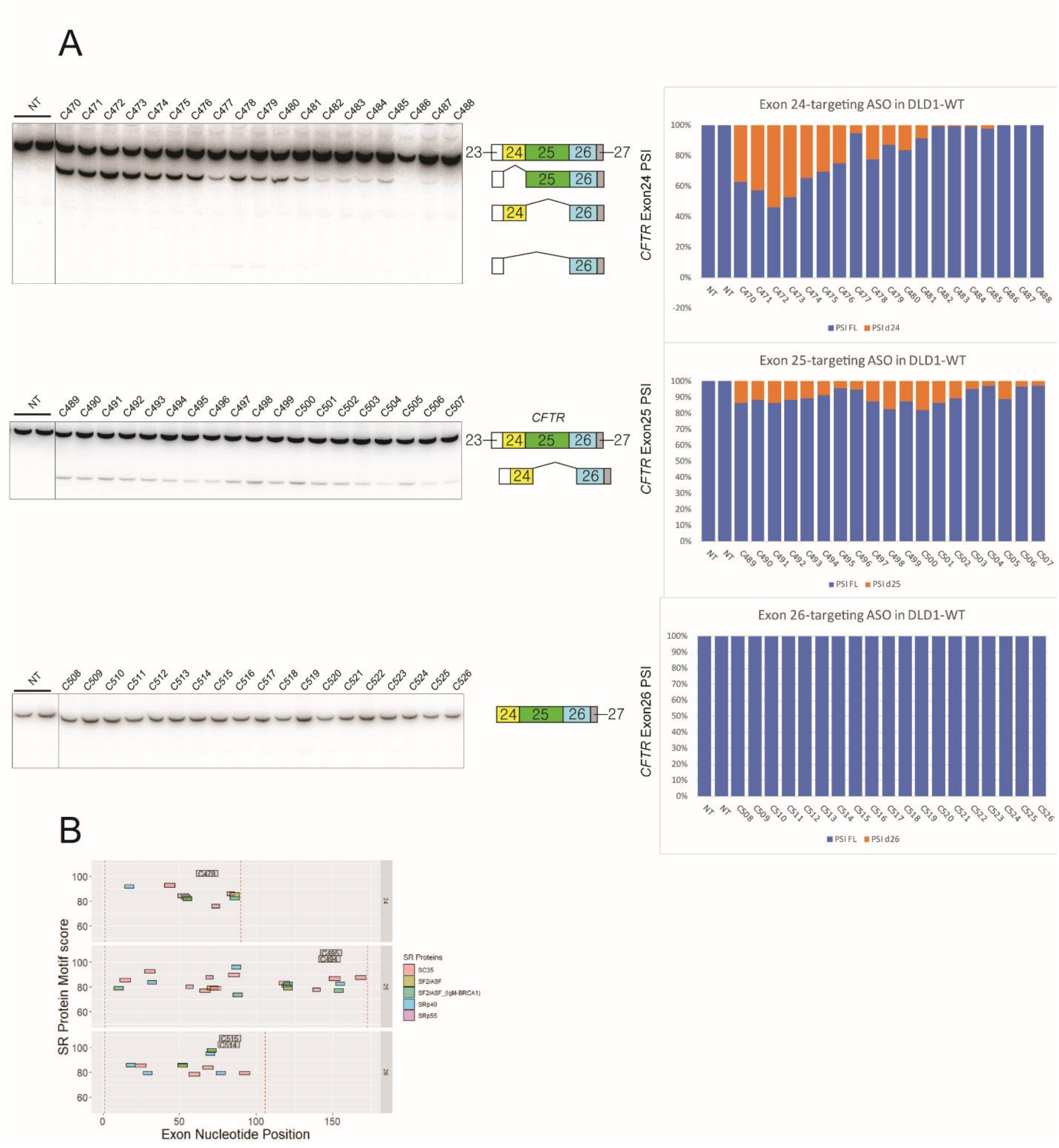


**Supplementary Figure 7. Effects of single 15mer ASOs on splicing of *CFTR-WT* mRNA.**

**A.** All 57 candidate ASOs (19 ASOs per exon) were individually transfected at a nominal concentration of 50 nM into DLD1-WT cells. *CFTR* mRNA exon 24, 25, and 26 percent-spliced-in (PSI) values were calculated from the RT-PCR data. **B.** SR protein motif analysis by

ESEfinder. (Original SR protein names; new names: SC35 = SRSF2, SF2/ASF = SRSF1, SRp40 = SRSF5, SRp55 = SRSF6). The small grey boxes show the target sites of the indicated lead ASOs. NT=No treatment.

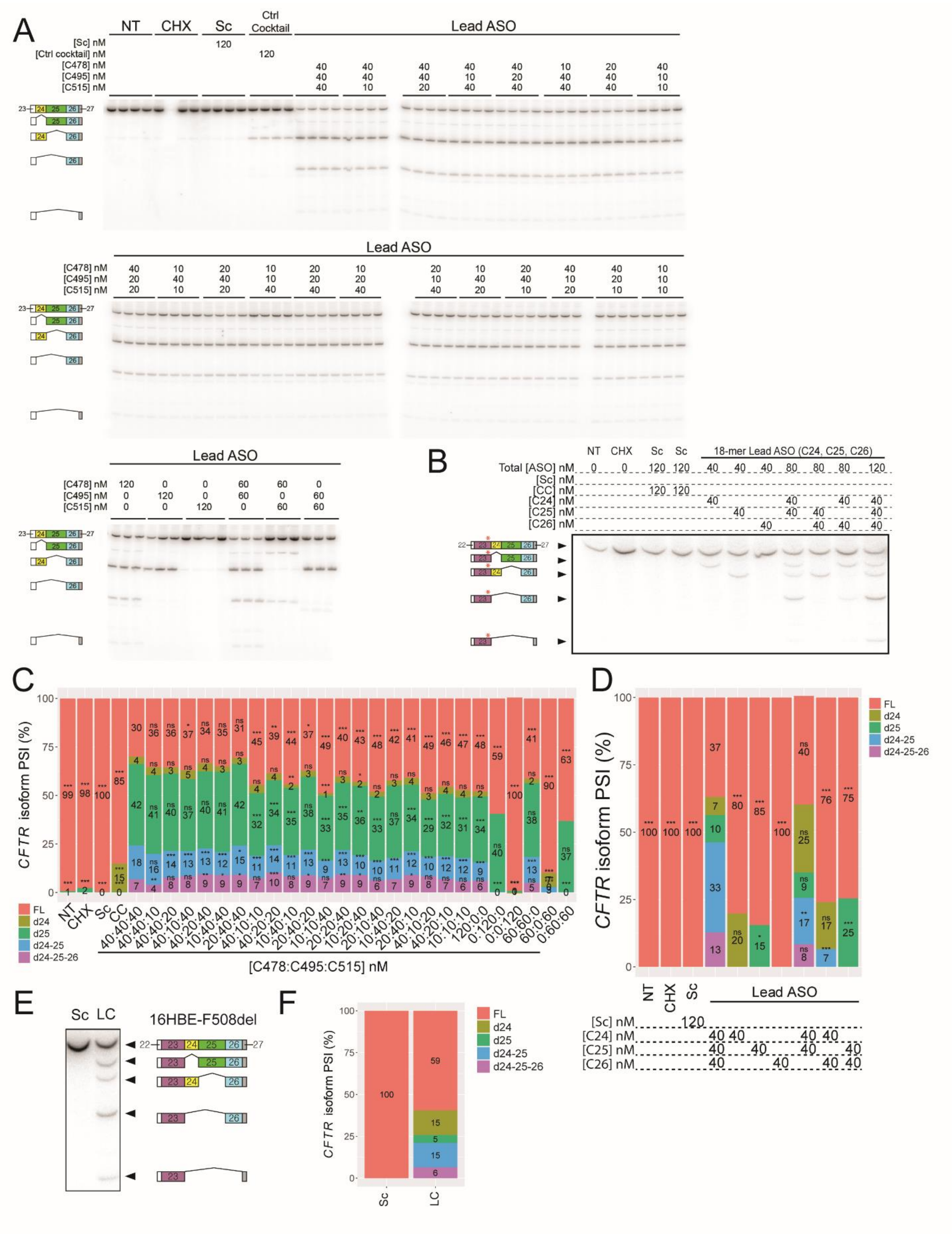


**Supplementary Figure 8. Effects of 15-mer or 18-mer ASO cocktails on *CFTR* splicing in**

**16HBE-W1282X cells; related to Figure 2H.**

**A.** Complete RT-PCR results of *CFTR* mRNA splicing patterns in 16HBE-W1282X cells transfected with various combinations of C478, C495, and C515 ASOs. **B.** Representative RT-PCR results of *CFTR* mRNA splicing patterns in 16HBE-W1282X cells transfected with various combinations of 18mer ASOs C24, C25, and C26. **C-D.** Mean PSI of each *CFTR* isoform in 16HBE-W1282X cells in (C) Panel A and (D) Panel B. **E.** Representative RT-PCR results of *CFTR* mRNA splicing patterns in 16HBE-F508del cells transfected with the lead 18mer ASO cocktail. **F.** Mean PSI of each *CFTR* isoform in 16HBE-F508del cells in Panel E. For Panel C-D, n=4 and 3, respectively. In Panels C-D, n.s. $P>0.05$, *$P<0.05$, **$P<0.01$, ***$P<0.001$, one-way ANOVA with Dunnett's post-test, versus each isoform in (Panel C) C478:C495:C515=40:40:40 nM or (Panel D) C24:C25:C26=40:40:40 nM. NT=No treatment; Sc=Scramble ASO; CC=Control ASO cocktail; CHX=cycloheximide. LC=Lead ASO cocktail C24-C25-C26.

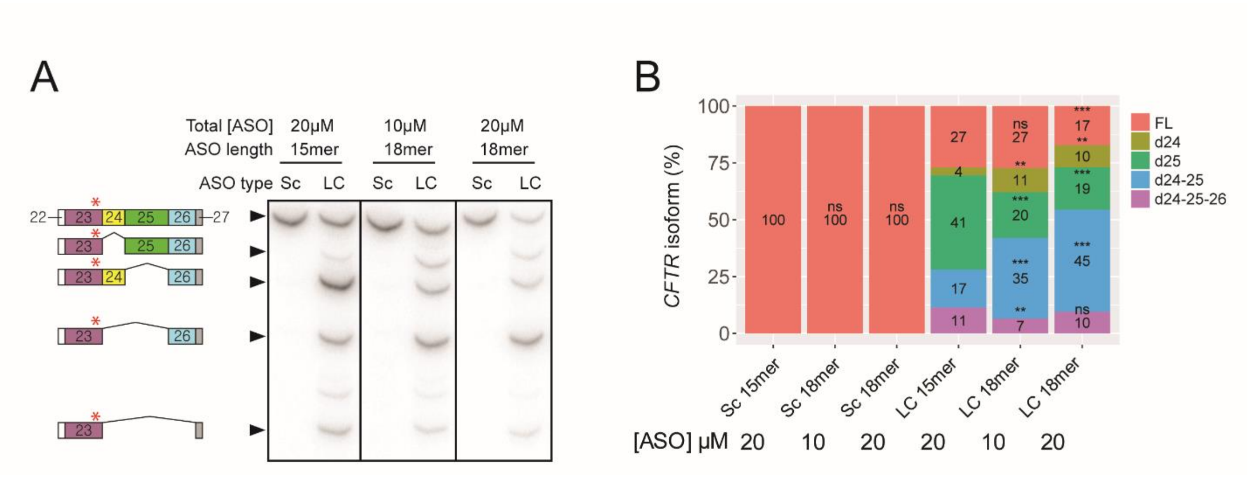


**Supplementary Figure 9. *CFTR* splicing in 16HBE-W1282X cells treated with 15mer or 18mer ASO cocktails by free-uptake.**

**A.** Representative RT-PCR results of *CFTR* mRNA splicing patterns in 16HBE-W1282X cells treated with 15mer lead cocktail (C478-C494-C515) or 18mer lead cocktail (C24-C25-C26). **B.** Mean PSI of each *CFTR* isoform in (A) (n=3 independent transfections, n.s. P>0.05, **P<0.01, ***P<0.001, one-way ANOVA with Dunnett's post-test, versus each isoform in Sc 15mer or LC 15mer). Sc=Scramble ASO; LC=Lead ASO cocktail C478-C494-C515 (15mer) or C24-C25-C26 (18mer).

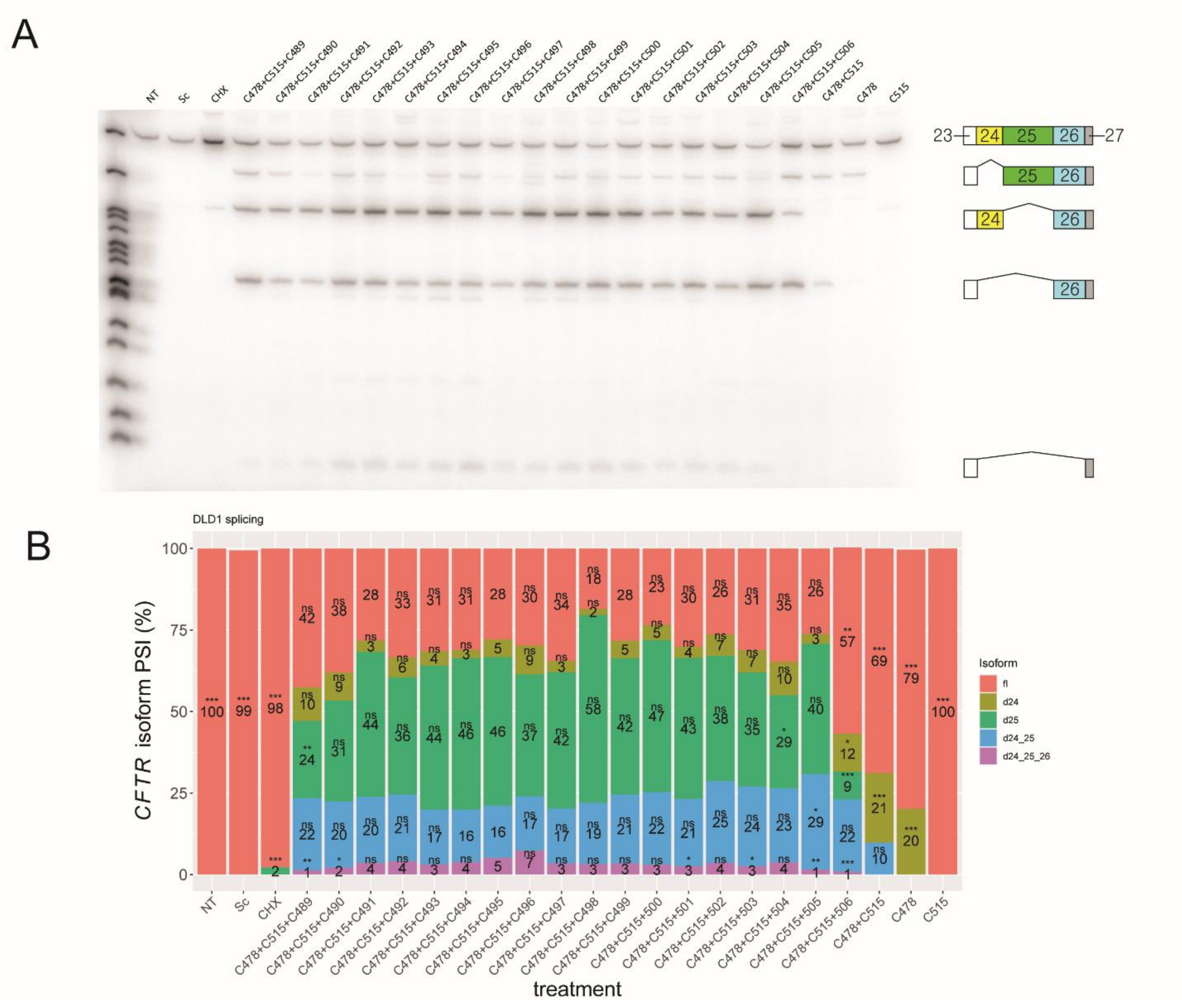

**Supplementary Figure 10. *CFTR* splicing in DLD1-W1282X cells transfected with 15mer ASO cocktails.**

**A.** Representative RT-PCR results of *CFTR* mRNA splicing patterns in DLD1-W1282X cells treated with various 15mer lead cocktails. **B.** Mean PSI of each *CFTR* isoform in DLD1-W1282X cells in (A) (n=3 independent transfections, n.s. P>0.05, *P<0.05, **P<0.01, ***P<0.001, one-way ANOVA with Dunnett's post-test, versus each isoform in C478:C495:C515=40:40:40 nM). NT= No treatment; Sc=Scramble ASO; CHX=cycloheximide.

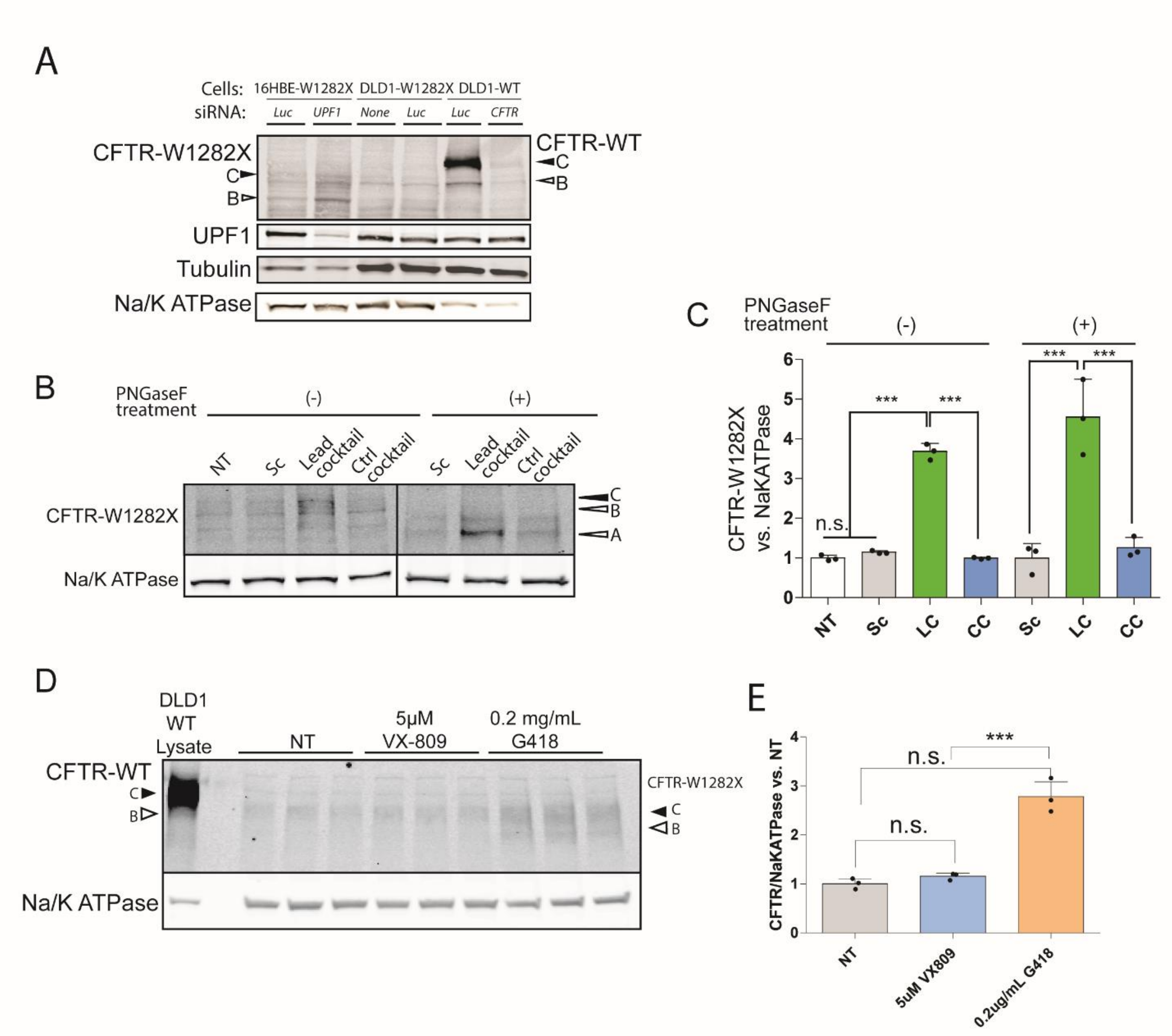


**Supplementary Figure 11. The EJC-targeting ASO cocktail increases CFTR-W1282X protein.**

**A.** CFTR antibody detects C and B bands of CFTR-WT and CFTR-W1282X. **B.** Western blot of complex glycosylated (C band), core-glycosylated (B-band), and non-glycosylated (A-band) of CFTR in 16HBE-W1282X cells transfected with 120 nM lead ASO cocktail (C478-494-C515). PNGase F was used to deglycosylate proteins in the cell extracts. Scramble 15-mer ASO (Sc) and control ASO cocktail (Ctrl Cocktail: C488-C507-C526) were used as negative controls. **C.** Western blot of 16HBE-W1282X cells treated with VX-809 or G418. **D.** Quantification of total

CFTR and deglycosylated CFTR protein in (B). (n=3 independent transfections, n.s. P>0.05, ***P<0.001, versus lead ASO cocktail, one-way ANOVA with Tukey's post-test). **E.** Quantification of CFTR-W1282X proteins in (C). (n=3 independent treatments, n.s. P>0.05, ***P<0.001, versus lead ASO cocktail, one-way ANOVA with Tukey's post-test). $Na^+/K^+$ ATPase served as internal reference. Western blot images are cropped images from the same SDS-PAGE gel.
